# A neuropeptide modulates sensory perception in the entomopathogenic nematode *Steinernema carpocapsae*

**DOI:** 10.1101/061101

**Authors:** Robert Morris, Leonie Wilson, Matthew Sturrock, Neil D. Warnock, Daniel Carrizo, Deborah Cox, Kilian McGrath, Aaron G. Maule, Johnathan J. Dalzell

## Abstract

Entomopathogenic nematodes (EPNs) employ a sophisticated chemosensory apparatus to detect potential hosts. Understanding the molecular basis of relevant host-finding behaviours could facilitate improved EPN biocontrol approaches, and could lend insight to similar behaviours in economically important mammalian parasites. FMRFamide-like peptides are enriched and conserved across the Phylum Nematoda, and have been linked with motor and sensory function, including dispersal and aggregating behaviours in the free living nematode *Caenorhabditis elegans*. The RNA interference (RNAi) pathway of *Steinernema carpocapsae* was characterised *in silico*, and employed to knockdown the expression of the FMRFamide-like protein 21 (GLGPRPLRFamide) gene (*flp-21*) in *S*. *carpocapsae* infective juveniles; a first instance of RNAi in this genus, and a first in an infective juvenile of any EPN species. Our data show that 5 mg/ml dsRNA and 50 mM serotonin triggers statistically significant *flp-21* knockdown (-84%***) over a 48 h timecourse, which inhibits host-finding (chemosensory), dispersal, hyperactive nictation and jumping behaviours. However, whilst 1 mg/ml dsRNA and 50 mM serotonin also triggers statistically significant *flp-21* knockdown (-51%**) over a 48 h timecourse, it does not trigger the null sensory phenotypes; statistically significant target knockdown can still lead to false negative results, necessitating appropriate experimental design. SPME GC-MS volatile profiles of two EPN hosts, *Galleria mellonella* and *Tenebrio molitor* reveal an array of shared and unique compounds; these differences had no impact on null *flp-21* RNAi phenotypes for the behaviours assayed. Localisation of *flp-21* / FLP-21 to paired anterior neurons by whole mount *in situ* hybridisation and immunocytochemistry corroborates the RNAi data, further suggesting a role in sensory modulation. These data can underpin efforts to study these behaviours in other economically important parasites, and could facilitate molecular approaches to EPN strain improvement for biocontrol.

**Author summary:** Entomopathogenic nematodes (EPNs) use a range of behaviours in order to find a suitable host, some of which are shared with important mammalian parasites. The ethical burden of conducting research on parasites which require a mammalian host has driven a move towards appropriate ‘model’ parasites, like EPNs, which have short life cycles, can be cultured in insects or agar plates, and have excellent genomic resources. This study aimed to develop a method for triggering gene knockdown by RNA interference, a biochemical pathway involved in gene regulation. Through knocking down the expression of a target gene we can then study the function of that gene, helping us to understand the molecular basis of behaviour. Here we have characterised the RNAi pathway of *Steinernema carpocapsae* through analysing the genome sequence for relevant genes, and have successfully knocked down the neuropeptide gene *flp-21* in *S*. *carpocapsae* infective juveniles. We find that it is involved in the regulation of behaviours which rely on sensory perception and relate to host-finding. We have localised the gene and mature neuropeptide, and find them to be expressed in paired anterior neurons, which is in broad agreement with our behavioural observations following RNAi. Our observations are relevant to interactions of *S*. *carpocapsae* with two insect hosts, the waxworm *Galleria mellonella*, and the meelworm, *Tenebrio molitor*. We identified the volatile compounds relating to both insects, and find that there are both shared and unique compounds to both species; EPNs use volatile compound gradients, as well as other physical cues in order to find and invade a host. This study provides a method for employing RNAi in a promising model parasite, and characterises the molecular basis of host-finding behaviours which could be relevant to economically important mammalian parasites. EPNs are also used as bioinsecticides, and so understanding their behaviour and biology could have broad benefits across industry and academia.

## Introduction

Entomopathogenic nematodes (EPNs) borrow their name from the entomopathogenic bacteria (*Photorhabdus*, *Serratia* and *Xenorhabdus* spp.) with which they form a commensal relationship. These nematodes provide a stable environment for the bacteria, and act as a vector between insect hosts. Once the nematode has invaded an insect, the nematode exsheaths (or ‘recovers’) and entomopathogenic bacteria are regurgitated into the insect haemolymph; the bacteria then rapidly kill and metabolise the insect, providing nutrition and developmental cues for the nematode. These entomopathogenic bacteria are then transmitted between nematode generations (1). The entomopathogenic lifestyle has been found to arise independently in nematodes, at least three times, spanning significant phylogenetic diversity. *Heterorhabditis* and *Oscheius* spp. (2) reside within clade 9 along with major strongylid parasites of man and animal (3); Steinernema spp. reside within clade 10 alongside strongyloidid parasites (4).

Nictation is a dispersal and host-finding strategy, enacted by nematodes which stand upright on their tails, waving their anterior in the air (5). This behaviour is shared amongst many economically important animal parasitic and entomopathogenic nematodes, alongside the model nematode *C. elegans*, for which nictation is a phoretic dispersal behaviour of dauer larvae, used to increase the likelihood of attachment to passing animals. Nictation is regulated by amphidial IL2 neurons in *C. elegans*, which occur in lateral triplets either side of the pharyngeal metacorpus (5, 6). IL2 neurons display significant remodelling from *C. elegans* L3 to dauer (the only life-stage to enact nictation behaviours) such that connectivity with other chemosensory and cephalic neurons is enhanced (6). It has been shown that IL2 neurons express the DES-2 acetylcholine receptor subunit, and that cholinergic signalling is requisite for nictation (5, 7–9). Additionally, the central pair of IL2 neurons express the FMRFamide-like peptide (FLP) receptor, NPR-1 (10). To date there are two known NPR-1 agonists; FLP-18 and FLP-21 (11). However, there is also known redundancy of FLP-18 and FLP-21 in signalling through other neuropeptide receptors (NPR-4, -5, -6 -10, -11, and NPR-2, -3, -5, -6, 11, respectively) in heterologous systems (12, 13), making functional linkage difficult. *Steinernema* spp. also display a highly specialised jumping behaviour which is thought to enhance both dispersal and host attachment. Jumping occurs when a nictating infective juvenile (IJ) unilaterally contracts body wall muscles bringing the anterior region towards the posterior region, forming a loop. This generates high pressure within the IJ pseudocoel, and differential stretching and compression forces across the nematode cuticle. Release of this unilateral contraction, in conjunction with the correction of cuticle pressure, triggers enough momentum for an IJ to jump a distance of nine times body length, to a height of seven times body length (14). Here we aimed to study the function of *Sc-flp-21* in coordinating nictation and other behaviours relevant to host-finding.

The recent publication of five *Steinernema* spp. genomes, along with stage-specific transcriptomes (15) represents a valuable resource, alongside the previously published genomes of *Oscheius* sp. TEL-2014 (16) and *Heterorhabditis bacteriophora* (17). The genome of *Steinernema carpocapsae* is the most complete, at an estimated 85.6 Mb, with predicted coverage of 98% (15). *S*. *carpocapsae* was selected as a test subject for our study due to the quality of genome sequence. The close phylogenetic relationship between *Steinernema* spp. coupled with a diverse behavioural repertoire, particularly in terms of host-finding (18, 19), make this genus an extremely attractive model for comparative neurobiology. The aim of this study was to examine RNAi functionality in *S*. *carpocapsae* IJs, and to probe the involvement of FLP-21 in coordinating sensory perception (host-finding, nictation, jumping and dispersal phenotypes), as a prelim to probing the neuronal and molecular underpinnings of host-finding behaviour in this genus.

## Materials and Methods

### *S*. *carpocapsae* culture

*S*. *carpocapsae* (ALL) was maintained in *Galleria mellonella* at 23°C. IJs were collected by White trap (20) in a solution of Phosphate Buffered Saline (PBS). Freshly emerged IJs were used for each experiment.

### BLAST analysis of *S*. *carpocapsae* RNAi pathway

BLAST analysis of RNAi pathway components was conducted as in Dalzell et al. (21), using a modified list of core RNAi pathway components from *C. elegans*, against predicted protein sets and contigs of the *S*. *carpocapsae* genome, through the Wormbase Parasite BLAST server (22, 23).

### dsRNA synthesis

*Sc-flp-21* (Gene ID: L596_g19959.t1) dsRNA templates were generated from *S*. *carpocapsae* IJ cDNA using gene-specific primers with T7 recognition sites (see Table 1). Neomycin phosphotransferase (*neo*) and Green Fluorescent Protein (*gfp*) dsRNA templates were generated from pEGFP-N1 (GenBank: U55762.1). Template PCR products were generated as follows: [95 °C x 10 min, 40 x (95 °C x 30 s, 60 °C x 30 s, 72 °C x 30 s) 72 °C x 10 min]. PCR products were assessed by gel electrophoresis, and cleaned using the Chargeswitch PCR clean-up kit (Life Technologies). dsRNA was synthesised using the T7 RiboMAX™ Express Large Scale RNA Production System (Promega), and quantified by a Nanodrop 1000 spectrophotometer.

### RNAi

1000 *S*. *carpocapsae* were incubated in 50 μl pBS with dsRNA and 50 mM serotonin across four experimental regimes; (i) 24 h in 5 mg/ml dsRNA / serotonin / PBS; (ii) 24 h in 5 mg/ml dsRNA / serotonin / PBS, followed by washes to remove the initial dsRNA, and 24 h recovery in PBS only; (iii) 48 h in 5 mg/ml dsRNA / serotonin / PBS; and (iv) 48 h in 1 mg/ml dsRNA and serotonin. Each experiment was conducted at 23 °C. We found that 50 mM serotonin induced oral uptake of fluorescent dyes under all conditions tested; 50 mM octopamine did not (data not shown).

### RNA extraction, cDNA synthesis and quantitative (q)RT-PCR

Total RNA was extracted from 1000 IJs using the Simply RNA extraction kit (Promega, UK) and Maxwell 16 extraction system (Promega, UK). cDNA was synthesised using the High Capacity RNA to cDNA kit (Applied Biosystems, UK). Each individual qRT-PCR reaction comprised 5 μl Faststart SYBR Green mastermix (Roche Applied Science), 1 μl each of the forward and reverse primers (10 μM), 1 μl water, 2 μl cDNA. PCR reactions were conducted in triplicate for each individual cDNA using a Rotorgene Q thermal cycler under the following conditions: [95 °C x 10 min, 40 x (95 °C x 20 s, 60 °C x 20 s, 72 °C x 20 s) 72 °C x 10 min]. Primer sets were optimised for working concentration, annealing temperature and analysed by dissociation curve for contamination or non-specific amplification by primer–dimer as standard. The PCR efficiency of each specific amplicon was calculated using the Rotorgene Q software. Relative quantification of target transcript relative to two endogenous control genes (*Sc-act* and *Sc-β-tubulin*) was calculated by the augmented ΔΔCt method (24), relative to the geometric mean of endogenous references (25). One way ANOVA and Fisher’s LSD test were used to analyse data (GraphPad Prism 6). The most similar non-target gene (*L596_g5821.t1*) was identified using BLASTn against the *S*. *carpocapsae* genomic contigs (supplemental figure S1), and primers *Sc-L596_g5821.t1-f* and *Sc-L596_g5821.t1-r* were used to assess transcript abundance relative to *Sc-act* across control and experimental conditions for the 48h dsRNA exposure experiments only (Table 1).

**Table 1.**
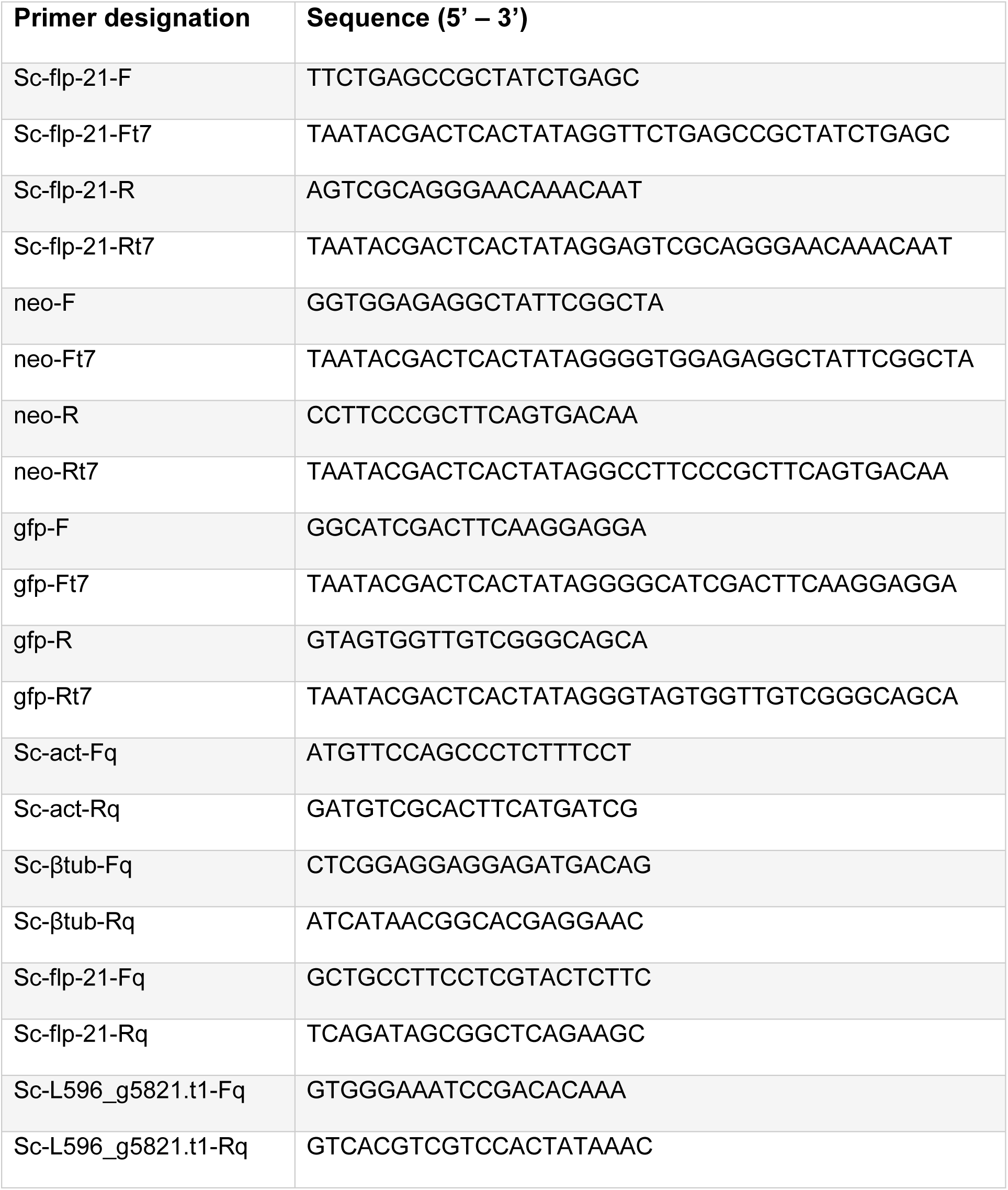
dsRNA synthesis and qRT-PCR primer sequences.

### Headspace Solid-Phase MicroExtraction (SPME) GC-MS

Approximately 5 g of fresh waxworm (*Galleria mellonella*) and mealworm (*Tenebrio molitor*) larvae were placed into 20 mL glass tubes and sealed. The holder needle was exposed to the headspace of the tube over a 120 min timecourse (extraction time) at room temperature (22 °C). After this time, the SPME syringe was directly desorbed in the GC injection port for 5 min. A fused silica fibre coated with a 95 μm layer of carboxen–polydimethylsiloxane (CAR–PDMS; Supelco) was used to extract the volatile compounds from the samples. Fibres were immediately thermally desorbed in the GC injector for 5 min (with this time we desorb the analytes and re-activated the fiber for the next analysis) at 250 °C and the compounds were analysed by GC-MS.

A CTC Analytics CombiPal autosampler was coupled to a 7890N Agilent gas chromatograph (Agilent, Palo Alto, California) and connected to a 5975C MSD mass spectrometer. The manual SPME holder (Supelco, Bellefonte, PA, USA) was used to perform the experiments. Chromatographic separation was carried out on 30 m x 0.25 mm I.D. ZB-semivolatiles, Zebron column (Phenomenex, Macclescfield, UK).The oven temperature was set at 40 °C for 3 min, temperature increased from 40 to 250 ° C at 5 °C min^−1^ and set at the maximum temperature for 4 min. Helium was used as carrier gas at 1 ml min^−1^. Mass spectra were recorded in electron impact (EI) mode at 70 eV. Scan mode was used for the acquisition to get all the volatile compounds sampled. Quadrupole and source temperature were set at 150 and 230 °C respectively. Compounds were identified using MS data from the NIST library (>95% confidence).

### Dispersal assays

100 *S*. *carpocapsae* IJs were placed in the centre of a 90 mm PBS agar plate (1.5 % w/v) in a 5 μl aliquot of PBS. Plates were divided into four zones; a central zone 15 mm in diameter, and three further zones equally spaced over the remainder of the plate. Plates were allowed to air dry for ~5 min. Evaporation of the PBS allowed the IJs to begin movement over the agar surface. Lids were then placed back onto the Petri dishes, and plates were incubated at 23 °C in darkness for one hour. IJs were counted across central and peripheral zones and expressed as percentage of total worms. Our subsequent analysis was conducted on total IJs found within the two central zones. Relative to those found in the two peripheral zones.

### Nictation & jumping assays

3.5 g of compost (John Innes No.2) was placed in a petri dish (55 mm), and dampened evenly with 150 μl PBS. Approximately ten IJs were pipetted onto the compost in 5 μl of PBS, and left for 5 minutes at room temperature; this enabled IJs to begin nictating. For the waxworm volatile challenge, one healthy waxworm (UK Waxworms Ltd.) was placed inside a 1 mL pipette tip, without filter. For the mealworm volatile challenge, two mealworms (Monkfield Nutrition, UK), weight-matched to the waxworm, were placed inside a 1 mL pipette tip, without filter. Blank exposure data were captured using an empty 1 ml pipette tip, without filter. In each case, the pipette was set to eject a volume of 500 μL, comprising air and the corresponding insect volatiles. A binocular microscope was used to record IJ behavioural responses following up to five volatile exposures each, on gentle ejection from the pipette within a distance of ~1 cm of the *S*. *carpocapsae* IJs. A five second period was allowed between each volatile exposure. Recording ended for any individual when jumping was observed or the IJ abandoned a nictating stance (this always corresponded with migration away from the stimulus). A jumping index was calculated for each treatment group (1). Additional behavioural observations were recorded, and subsequently reported as percentage IJs displaying the behaviour over the course of up to five volatile exposures, or until the IJ migrated / jumped out of the field of vision.

### Host finding assay

Two circular holes (approx. 6 mm diameter, centred 4 mm from edge of lid) were drilled either side of a 90 mm petri dish lid. Two microcentrifuge tubes (1.5 ml) with a small hole cut out the bottom (approx. 2mm diameter), were also used for each assay. 200 *S*. *carpocapsae* IJs were placed in the centre of a 90 mm PBS agar plate (1.5 % w/v) in a 5 μl aliquot of PBS. The arena was segmented into positive and a negative zones either side of the plate (25 mm in length from the edge, circling off the plate at a point 60 mm apart; see Fig XA). Plates were allowed to air dry for ~5 min, allowing the IJs to begin migration. The lid was placed on top of the plate, and sealed with parafilm. The 1.5 ml tubes were secured in the holes with parafilm; one remained empty, which we term the blank tube, and the other held four live *Galleria mellonella* fourth instar larvae, or four *Tenebrio molitor* larvae as appropriate. The lid of the tubes were then closed. The plates were incubated at 23 °C in darkness for one hour. IJs were counted in the positive (host side) and negative (blank side) zones and then scored using a chemotaxis index (26). The assay format was adapted from Grewal et al. (1994) (27).

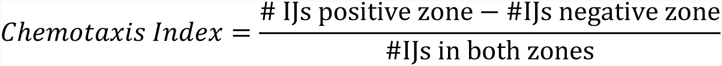

### Immunocytochemistry

Freshly emerged *S*. *carpocapsae* IJs were fixed in 4 % paraformaldehyde overnight at 4 °C, followed by a brief wash in antibody diluent (AbD; 0.1 % bovine serum albumin, 0.1 % sodium azide, 0.1 % Triton-X-100 and PBS pH 7.4). The fixed specimens were roughly chopped on a glass microscope slide with a flat edged razor, and incubated in primary polyclonal antiserum raised against GLGPRPLRFamide, N-terminally coupled to KLH, and affinity purified (1:800 dilution in AbD) for 72 h at 4 °C. Subsequently, chopped IJs were washed in AbD for 24 h at 4 °C, and then incubated in secondary antibody conjugated to fluorescein isothiocyanate (1:100 dilution in AbD) for 72 h at 4 °C. A further AbD wash for 24 h at 4 °C was followed by incubation in Phalloidin–Tetramethylrhodamine B isothiocyanate (1:100 dilution in AbD) for 24 h at 4 °C. Finally, chopped IJs were washed in AbD for 24 h at 4 °C. Specimens were mounted onto a glass slide with Vectasheild mounting medium and viewed with a Leica TCS SP5 confocal scanning laser microscope. Controls included the omission of primary antiserum, and pre-adsorption of the primary antiserum with ≥250 ng of GLGPRPLRFamide. Pre-adsorption in GLGPRPLRFamide did not alter staining patterns.

### Whole mount *in situ* hybridisation

PCR primers were designed to amplify a 200 bp region of *Sc-flp-21* (Gene ID: L596_g19959.t1) from *S. carpocapse* IJ cDNA. Template PCR products were generated as follows : [95°C x 10 min, 40 × (95 °C x 30 sec, 60 °C x 30 sec, 72 °C x 30 sec) 72 °C x 10 min]. PCR products were assessed by gel electrophoresis, and cleaned using the Chargeswitch PCR clean-up kit (Life Technologies). Amplicons were quantified by a Nanodrop 1000 spectrophotometer. Sense and antisense probes were generated using amplicons in an asymmetric PCR reaction. The components of each reaction were as follows: 2.0μl of Reverse primer (or Forward primer for control probe); 2.5μl 10x PCR buffer with MgCl2 (Roche Diagnostics); 2μl DIG DNA labelling mix (Roche Diagnostics); 0.25μl 10x *Taq* DNA polymerase (Roche Diagnostics); 20ng DNA template; distilled water to a volume of 25μl. Probes were assessed by gel electrophoresis.

Freshly emerged *S*. *carpocapse* IJs were fixed in 2 % paraformaldehyde in M9 buffer overnight at 4°C followed by 4h at room temperature. Nematodes were chopped roughly using a sterile razor blade for 2 minutes and washed three times in DEPC M9. Subsequently, the chopped nematodes were incubated in 0.4 mg/ml proteinase K (Roche Diagnostics) for 20 minutes at room temperature. Following three washes in DEPC M9, the nematodes were pelleted (7000g) and frozen for 15 minutes on dry ice. Subsequently the nematode sections were incubated for 1 minute in -20°C methanol and then in -20°C acetone for 1 minute. The nematodes were then rehydrated using DEPC M9 and incubated at room temperature for 20 minutes, after which three wash steps in DEPC M9 were carried out to remove any acetone.

The nematodes were pre-hybridised in 150 μl hybridisation buffer (prepared as detailed by Boer et al., 1998) for 15 minutes. The hybridisation probes were heat denatured at 95°C for 10 minutes, after which they were diluted with 125 μl hybridisation buffer. The probe-hybridisation mixture was then added to the nematode sections which were incubated at 50°C overnight. Post hybridisation washes were carried out as follows: three washes in 4x Saline Sodium Citrate buffer (15 minutes, 50°C); three washes in 0.1× SSC/0.1x Sodium dodecyl sulphate (20 minutes, 50°C) and; 30 minute incubation in 1% blocking reagent (Roche Diagnostics) in maleic acid buffer (50°C). Subsequently the nematodes were incubated at room temperature for 2 h in alkaline phosphatase conjugated anti-digoxigenin antibody (diluted 1:1000 in 1% blocking reagent in maleic acid buffer). Detection was completed with an overnight incubation in 5-Bromo-4-chloro-3-indolyl phosphate/Nitro blue tetrazolium at 4°C. The staining was stopped with two washes in DEPC treated water. The nematode sections were mounted on to glass slides for visualisation.

### Statistical analysis

Data pertaining to both qRT-PCR and behavioural assays were assessed by Brown-Forsythe and Bartlett’s tests to examine homogeneity of variance between groups. One-way or two-way ANOVA was followed by Fisher’s Least Significant Difference (LSD) test. All statistical tests were performed using GraphPad Prism 6.

## Results

### The RNAi pathway of *S*. *carpocapsae*

As is the case for other parasitic nematode species, *S. carpocapsae* was found to encode a less diverse RNAi pathway than that of *C. elegans*, in terms of gene for gene conservation (21). However, the apparent reduction in AGO homologue diversity is offset by significant expansions across several putative *ago* genes, to give a predicted overall increase in the *S. carpocapsae* AGO complement (38 in total), relative to *C. elegans* (24, not including pseudogenes) (28); WAGO-1 (nineteen), ALG-1 (three), ALG-3 (two), WAGO-5 (four), WAGO-10 (two), WAGO-11 (three) are all expanded relative to *C. elegans*. Notably, no identifiable homologue of RDE-1, the primary AGO for exogenously triggered RNAi events in *C. elegans*, could be identified (Supplementary table S2).

The presence of PRG-1 and components of the piwi interacting (pi)RNA biosynthetic machinery suggests that a functional piRNA (or 21U RNA) pathway may be present. Whilst ERGO-1 is not conserved, two putative ALG-3 orthologues suggest that a functional endo-siRNA (26G RNA) pathway may also exist, which is supported by broad conservation of associated proteins. MicroRNA-associated AGOs, ALG-1 and ALG-2 are conserved, with a small apparent expansion of ALG-1 to three related proteins in *S*. *carpocapsae*. Further understanding of how RNAi pathway complements influence functionality will require small RNA sequencing efforts, and functional genomics approaches.

The RNA-dependent RNA Polymerase (RdRp), RRF-3 is conserved, and known to function antagonistically to exogenously primed RNAi, through competing activity for pathway components required for both exogenous RNAi, and the endo-siRNA (26G RNA) pathway within which RRF-3 operates (29–31). The RdRps, RRF-1 and EGO-1, which are involved in the biosynthesis of secondary siRNAs (22G RNAs) are also conserved. Loss of the argonaute ERGO-1 which functions upstream of secondary siRNA biogenesis in the endo-siRNA (26G RNA) pathway in *C. elegans*, also leads to an exogenous ERI phenotype (Enhanced RNAi), but is not conserved in S, *carpocapsae*, suggesting that ALG-3 / -4 may be solely responsible for endo-siRNA functionality (32, 33).

The apparent absence of the intestinal dsRNA transporter, SID-2 is consistent with findings from other parasitic nematodes (21, 34, 35). SID-1 also appears to be absent, however CHUP-1, a putative cholesterol uptake protein which contains a SID-1 RNA channel is present, and may assist in the intercellular spread of dsRNA. RSD-3, which also effects intercellular spread of dsRNA is conserved (see Figure 1 for pathway overview).

**Figure 1.**
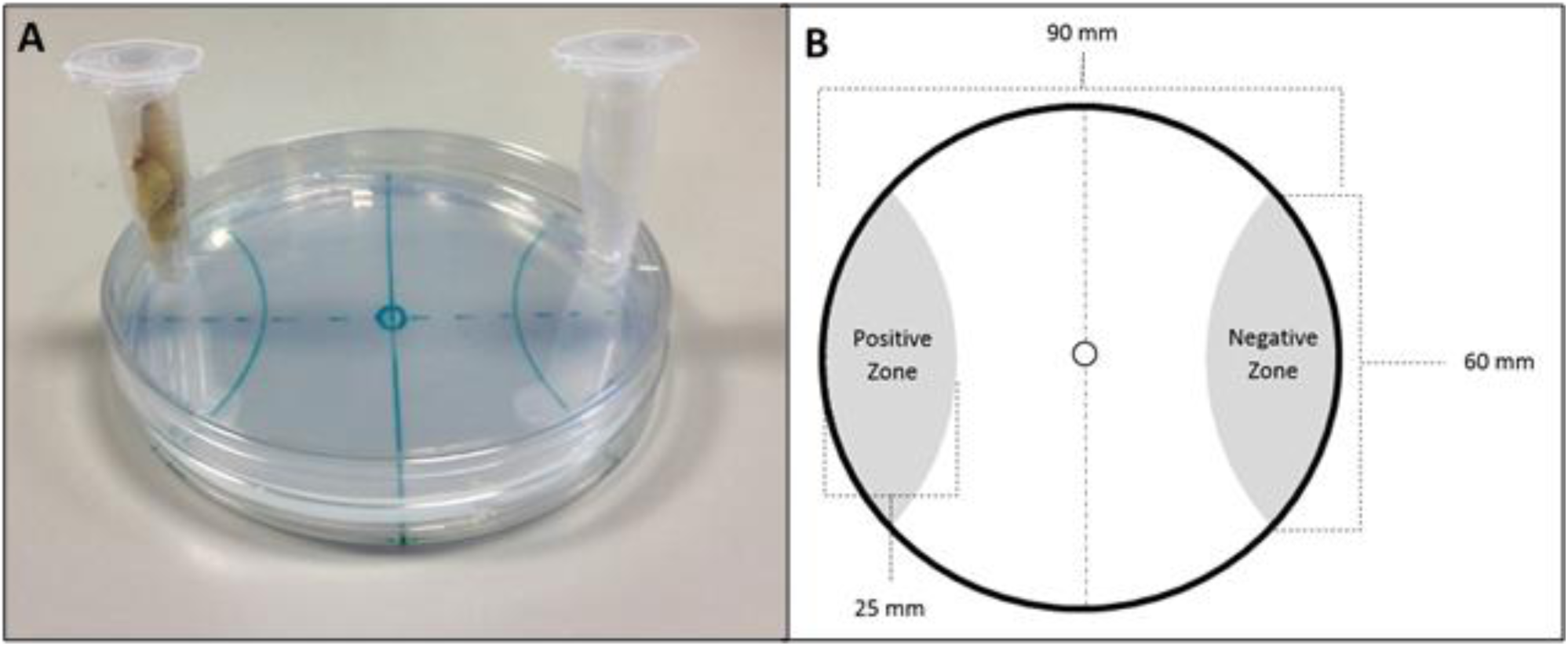
(A)image of the final host-finding assay; (B) host-finding assay schematic.

**Figure 1.**
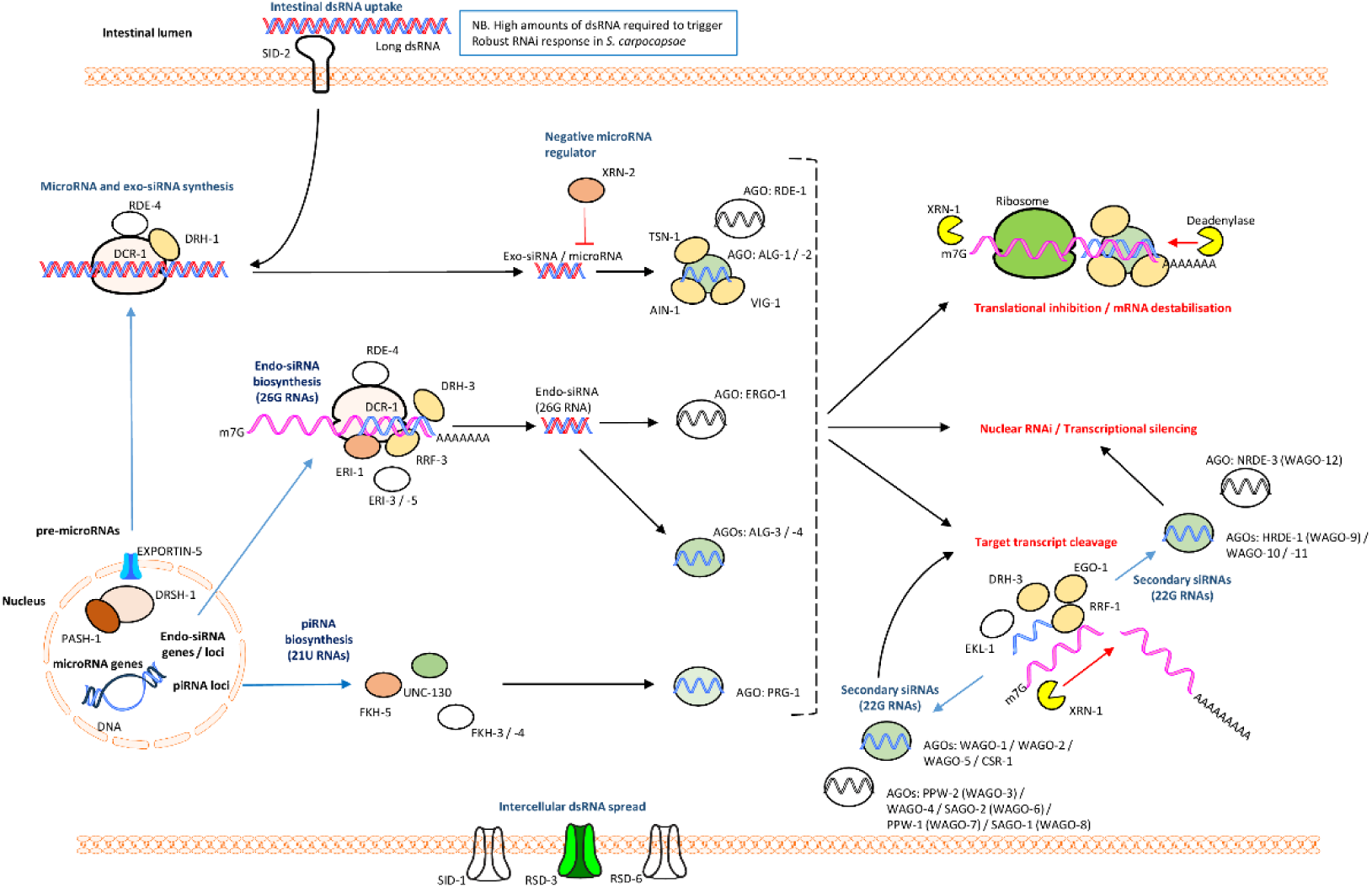
Core RNAi pathway components of *S*. *carpocapsae* relative to *C. elegans*.

Proteins with at least one putative homologue are represented in colour; those without any identifiable homologues are colourless. See supplemental tables for information on number of putative related homologues.

### Knockdown of *Sc-flp-21*

Various treatment regimens were employed in order to assess the responsiveness of *S*. *carpocapsae* IJs to exogenous dsRNA. 24 h incubation in 5 mg/ml dsRNA, with 50 mM serotonin was not sufficient to trigger statistically significant *Sc-flp-21* knockdown (Fig. 2A), however a 24 h dsRNA / serotonin incubation followed by a 24 h recovery in PBS only, did trigger a small decrease in *Sc-flp-21* relative to *Sc-act* when compared to *gfp* and *neo* dsRNA controls (0.70 ±0.11, P<0.05) (Fig. 2B). Extended incubation of *S*. *carpocapsae* IJs in 5 mg/ml dsRNA and 50 mM serotonin for 48 h triggered robust knockdown of *Sc-flp-21* (0.16 ±0.07, P<0.0001) (Fig. 2C). 48 h incubation in 1 mg/ml dsRNA, with 50 mM serotonin also triggered significant levels of *Sc-flp-21* knockdown (0.49 ±0.27, P<0.01), however this was not as effective as the 5 mg/ml dsRNA treatment (Fig. 2D). A BLAST analysis identified predicted *S*. *carpocapsae* transcript *L596_g5821.t* as the non-target gene with most similarity to the *Sc-flp-21* dsRNA (supplemental figure S1). The relative expression level of *L596_g5821.t1* was unaffected by a 48 h incubation in 5 mg/ml *Sc-flp-21* dsRNA with 50 mM serotonin, relative to *neo* and *gfp* dsRNA (1.013 ±0.04, P>0.05) (Fig. 2E).

**Figure 2.**
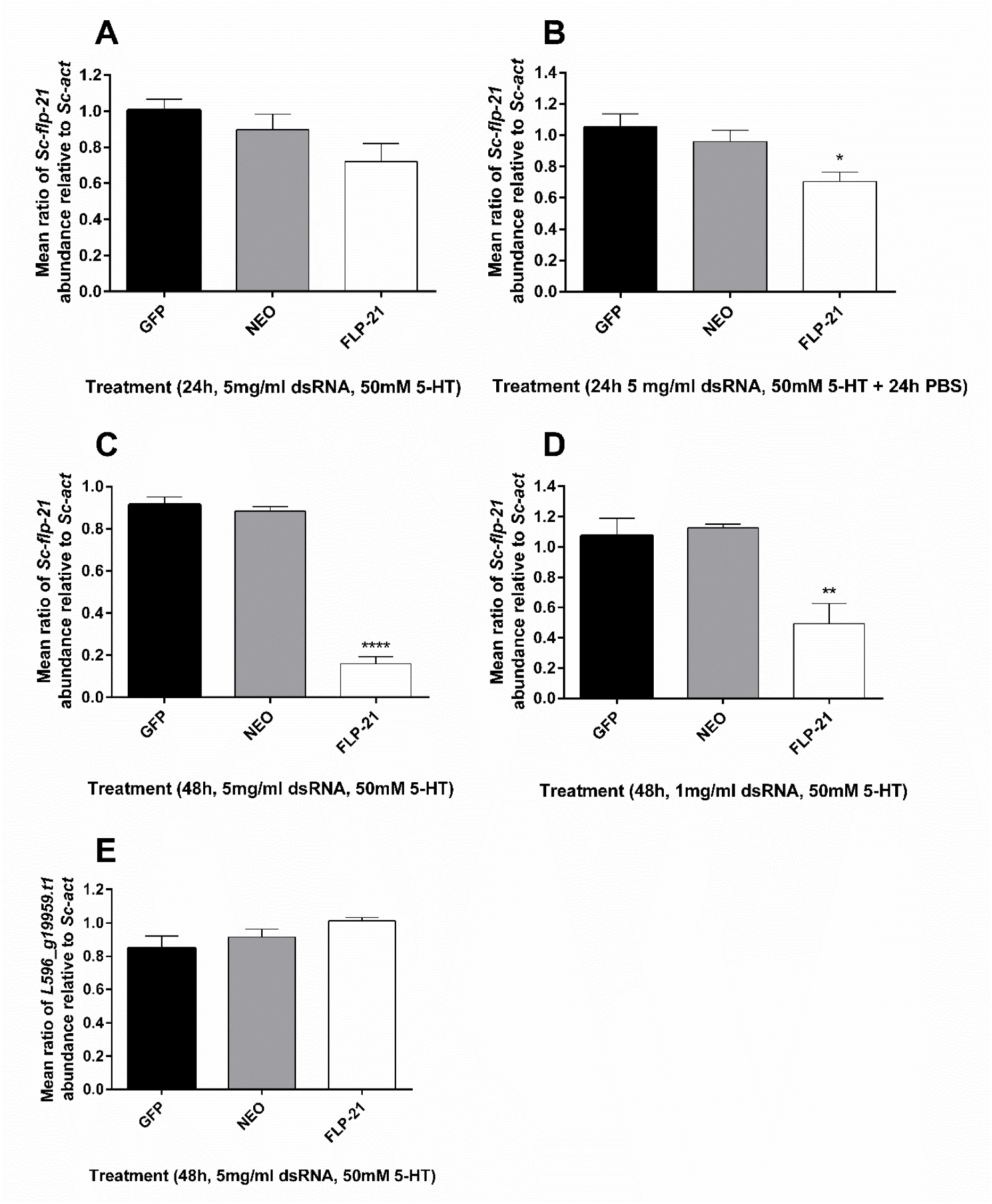
qRT-PCR expression analysis of *Sc-flp-21* and off-target control gene

(A)*Sc*-*flp-21* transcript ratio relative to *Sc-act* following 24 h incubation in 5 mg/ml dsRNA, 50 mM serotonin. (B) *Sc-flp-21* transcript ratio relative to *Sc-act* following 24 h incubation in 5 mg/ml dsRNA, 50 mM serotonin, and 24 h in PBS; (C) *Sc-flp-21* transcript ratio relative to *Sc-act* following 48 h incubation in 5 mg/ml dsRNA, 50 mM serotonin; (D) *Sc-flp-21* transcript ratio relative to *Sc-act* following 48 h incubation in a reduced 1 mg/ml dsRNA, 50 mM serotonin; (E) *L596_g5821.t1* transcript ratio relative to *Sc-act* following 48 h incubation in 5 mg/ml dsRNA, 50 mM serotonin. *P<0.05; **P<0.01; ****P<0.0001.

### Host insect volatiles

Comprehensive volatile signatures were characterised, and significant differences noted between *G. mellonella* and *T. molitor* larvae. In total, we identified 10 compounds unique to *G. mellonella*, four compounds unique to *T. molitor*, and 14 compounds shared between both species. These profiles vary significantly from headspace GC-MS data presented by Hallem et al. (36) for the same insect species (see Table 1).

**Table 1.**
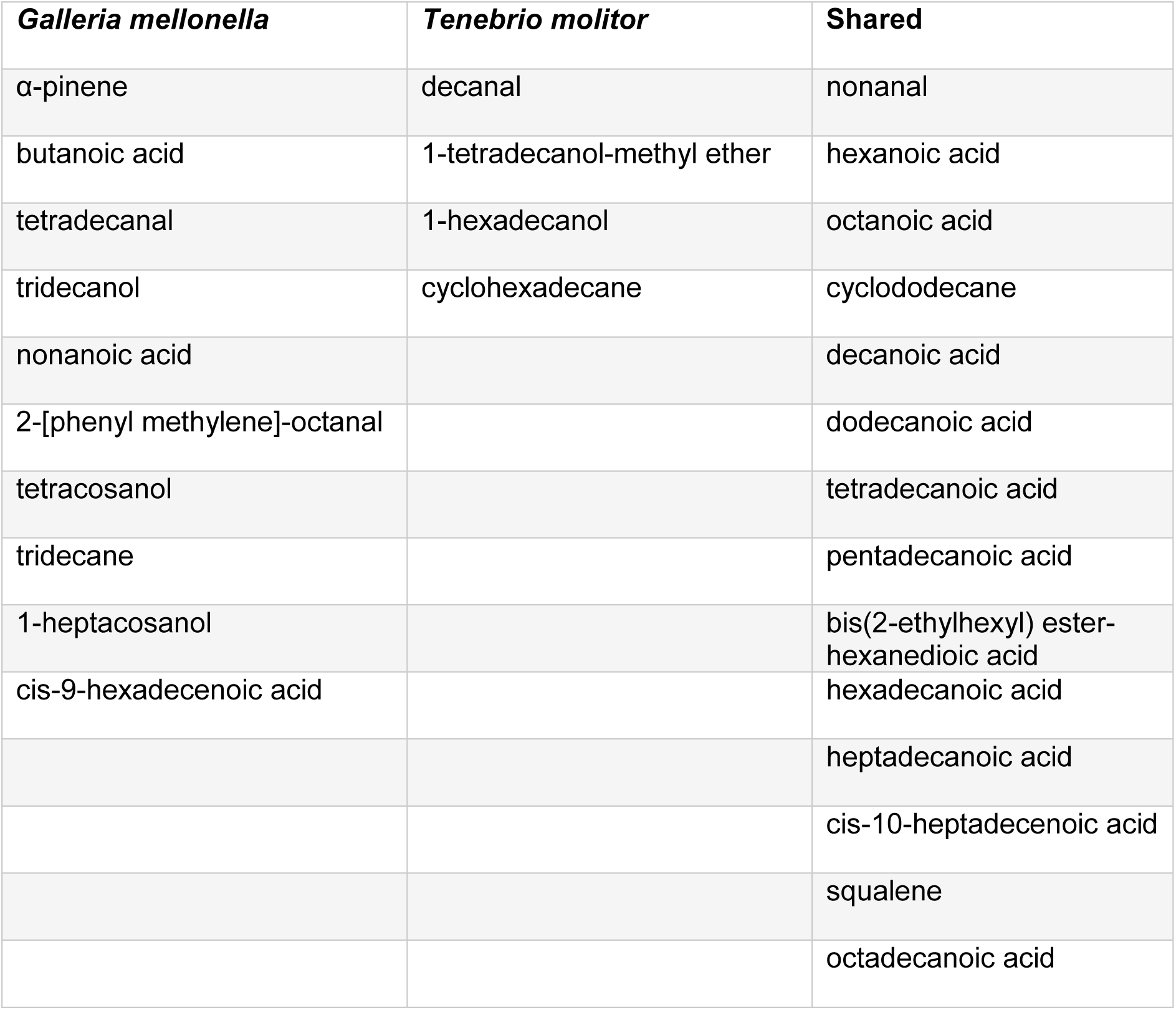
Headspace SPME GC-MS volatile profiles of *Galleria mellonella* and *Tenebrio molitor* larvae

| *Galleria mellonella* | *Tenebrio molitor* | Shared |
| --- | --- | --- |
| α-pinene | decanal | nonanal |
| butanoic acid | 1-tetradecanol-methyl ether | hexanoic acid |
| tetradecanal | 1-hexadecanol | octanoic acid |
| tridecanol | cyclohexadecane | cyclododecane |
| nonanoic acid |  | decanoic acid |
| 2-[phenyl methylene]-octanal |  | dodecanoic acid |
| tetracosanol |  | tetradecanoic acid |
| tridecane |  | pentadecanoic acid |
| 1-heptacosanol |  | bis(2-ethylhexyl) ester-hexanedioic acid |
| cis-9-hexadecenoic acid |  | hexadecanoic acid |
|  |  | heptadecanoic acid |
|  |  | cis-10-heptadecenoic acid |
|  |  | squalene |
|  |  | octadecanoic acid |

### Behavioural impact of *Sc-flp-21* knockdown

*S*. *carpocapsae* IJs were challenged by exposure to volatiles from *G. mellonella* or *T. molitor* following RNAi (48 h 5 mg/ml dsRNA, 50 mM serotonin) and control treatments. A decrease in hyperactive nictation following *Sc-flp-21* knockdown was observed (10% ±5.774) relative to untreated (40.75% ±6.75; P<0.01) and *neo* dsRNA treatment (47.5% ±2.5; P<0.001) following *G. mellonella* volatile challenge (Fig. 3A). Likewise, a decrease in hyperactive nictation was observed following *T. molitor* volatile challenge to *Sc-flp-21* RNAi IJs (5.0% ±2.9), relative to untreated (57.25% ±2.8; P<0.0001) and *neo* dsRNA treatment (35.0% ±6.5; P<0.001) (Fig. 3B). A decrease in the jumping index of IJs following *Sc.flp-21* dsRNA treatment was observed when challenged by *G. mellonella* volatiles (0.08 ±0.03) relative to untreated (0.69 ±0.11; P<0.001) and *neo* dsRNA treated (0.50 ±0.04; P<0.01) (Fig. 3E). Similarly, a decrease in jumping index as a response to *T. molitor* volatiles was observed following *Sc-flp-21* RNAi (0.03 ±0.03) relative to untreated (0.44 ±0.06; P<0.001) and *neo* dsRNA treatment (0.38 ±0.05; P<0.001) (Fig 3F). An agar host-finding assay was used to further assess the impact of *flp-21* knockdown. A decrease in *G. mellonella* finding ability was observed (0.06 ±0.08) relative to untreated (0.53 ±0.03; P<0.001) and *neo* dsRNA treated (0.42 ±0.08; P<0.01). Likewise, a decrease in *T. molitor* finding ability was observed (0.01 ±0.06) relative to untreated (0.32 ±0.04; P<0.05) and *neo* dsRNA treated (0.26 ±0.1; P>0.05). It was also found that *Sc-flp-21* RNAi resulted in significantly decreased lateral dispersal, relative to both untreated and *neo* dsRNA treatment (P<0.0001) (Fig. 3G). In all instances, dsRNA treatment regimens which triggered lower levels of *Sc-flp-21* knockdown relative to the 48h 5 mg/ml dsRNA, 50 mM serotonin approach, failed to trigger null phenotypes (data not shown).

**Figure 3.**
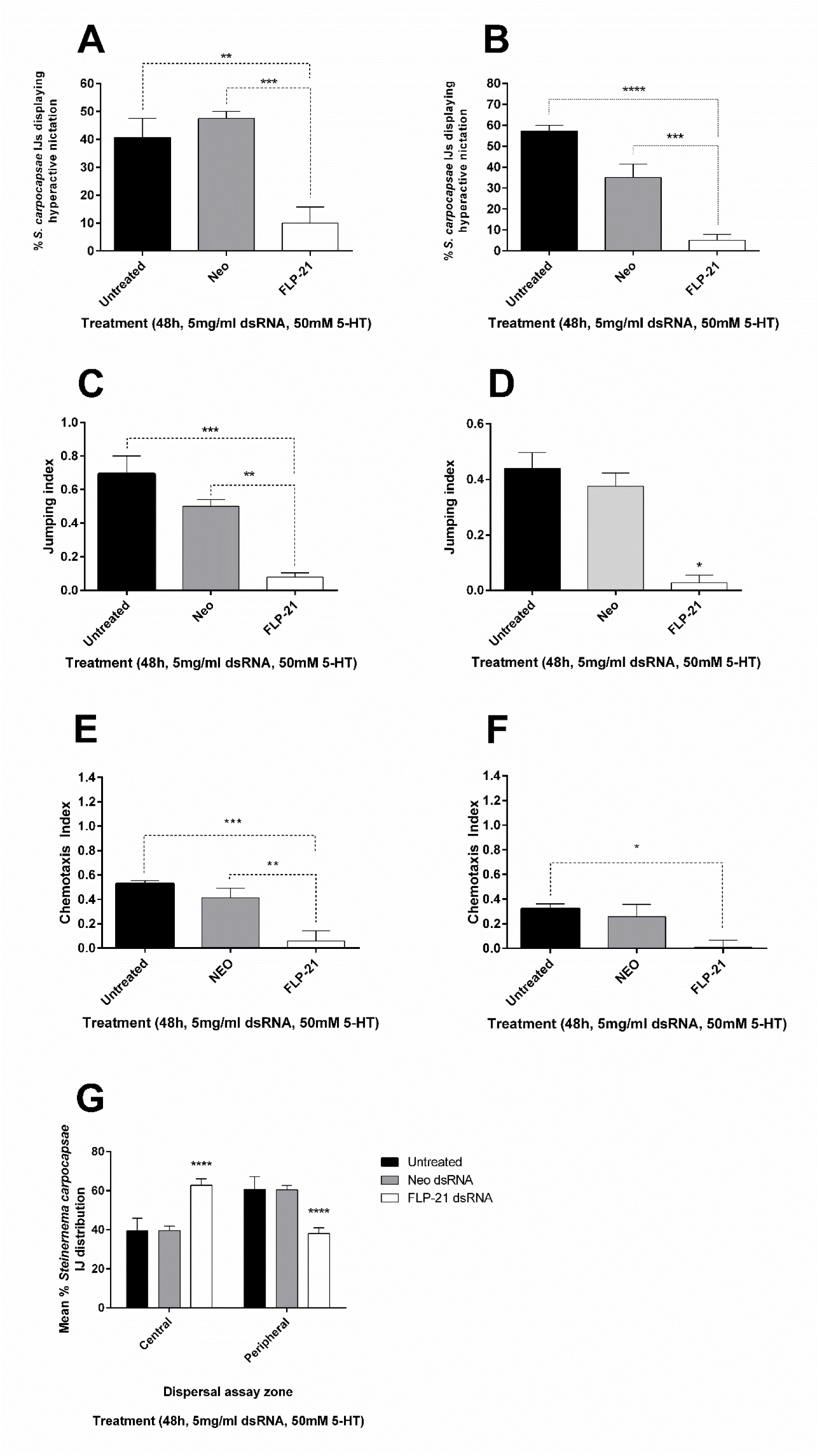
*S*. *carpocapsae* IJ behavioural assays post-RNAi

Behavioural impact of *Sc-flp-21* knockdown following IJ incubation in 5 mg/ml dsRNA, 50 mM serotonin: (A) Mean percentage of *S*. *carpocapsae* displaying hyperactive nictation upon challenge by *G. mellonella* volatiles. (B) Mean percentage of *S*. *carpocapsae* displaying hyperactive nictation upon challenge by *T. molitor* volatiles. (C) Mean percentage of *S*. *carpocapsae* displaying standing upon challenge by *G. mellonella* volatiles. (D) Mean percentage of *S*. *carpocapsae* displaying standing upon challenge by *T. molitor* volatiles. (E) Mean jumping index of *S*. *carpocapsae* displaying upon challenge by *G. mellonella* volatiles. (F)Mean jumping index of *S*. *carpocapsae* upon challenge by *T. molitor* volatiles. (G) Mean percentage distribution of *S*. *carpocapsae* IJs across central and peripheral assay zones. *P<0.05; **P<0.01; ****P<0.0001.

### Whole-mount *in situ* hybridisation and Immunocytochemical localisation of *flp-*21/FLP-21 in *S*. *carpocapsae* IJs

*flp-21*/FLP-21 was localised to paired neurons within the central nerve ring region of *S*. *carpocapsae* IJs. Without additional neuroanatomical information on *S*. *carpocapsae* IJs it is impossible to define these cells, however, based on the immuocytochemical localisation the cells appear to project posteriorly (Fig. 4). These data suggest that FLP-21 must act as a modulator of sensory function, downstream of the primary chemosensory neurons (amphids).

**Figure 4.**
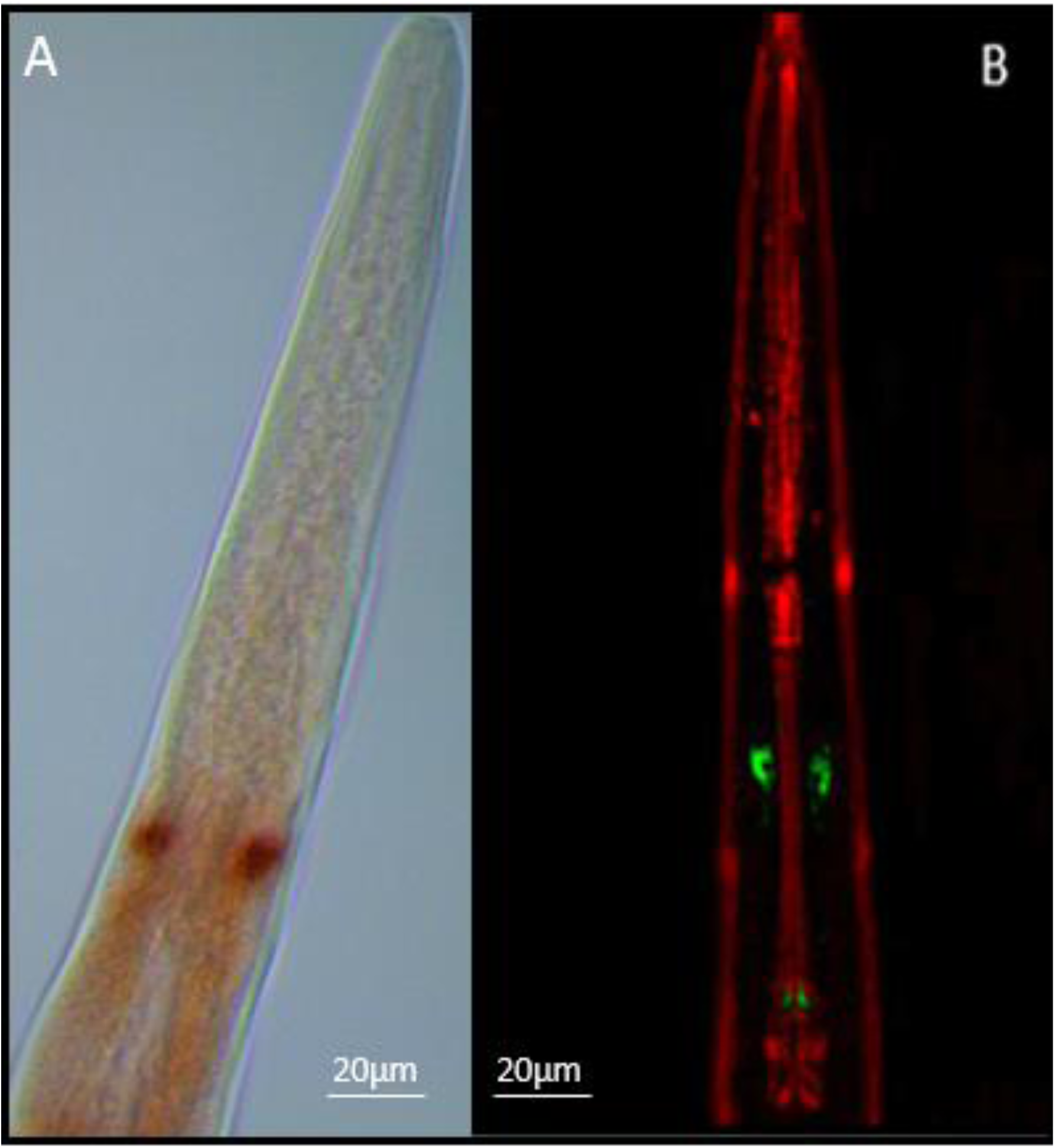
(A) Whole mount In situ hybridisation and (B) immunocytochemical localisation of *flp-21* / FLP-21 (GLGPRPLRFamide) to paired neurons within the central nerve ring of *S*. *carpocapsae* IJs.

Positive *flp-21* staining is observed as red/brown pigmentation (A); FLP-21 immunostaining is visible as green, muscle is counterstained red (B)

## Discussion

RNA interference is an extremely important tool for the study of gene function in parasitic nematodes (37, 38). Three independent reports of a functional RNAi pathway in the entomopathogenic nematode *Heterorhabditis bacteriophora* have been published. Ciche and Sternberg (39) assessed the efficacy of RNAi through soaking egg / L1 stage *H. bacteriophora* in 5-7.5 mg/ml dsRNA targeting a number of genes which had been selected on the basis of phenotypic impact on the model *C. elegans*. Demonstrable phenotypes and target transcript knockdown signified an active pathway. Moshayov, Koltai and Glazer (2013)(40) employed the methodology of Ciche and Sternberg (39)to study the involvement of genes in the regulation of IJ exsheathment (or ‘recovery’). Subsequently, Ratnappan et al. (41) demonstrated that microinjection was also a suitable method for introducing dsRNA into hermaphrodite gonads, effectively triggering the RNAi pathway in F1 progeny. To date, no such assessment of a functional RNAi pathway has been published for *Steinernema* spp.

The RNAi pathway of *S*. *carpocapsae* has been characterised by BLAST and validated through silencing *Sc-flp-21* in IJs. Our data indicate that neuronal cells are sensitive to RNAi in *S*. *carpocapsae* IJs, and that knockdown is highly sequence specific. Like other parasitic nematodes *S*. *carpocapsae* encodes an expanded set of WAGO-1 (R06C7.1) family AGOs (19 in total) which function primarily with secondary siRNAs (22G RNAs) in *C. elegans*, along with CSR-1 which is also conserved. Whilst RDE-1 is primarily responsible for triggering the onset of an exogenous RNAi response, acting upstream of secondary siRNAs (22G RNAs), it is not conserved in *S*. *carpocapsae* (21, 31). Our observation of RNAi sensitivity in *S. carpocapsae* reveals that RDE-1 is not required to trigger an exogenous RNAi response, however the functional significance of AGO homologue expansions relative to *C. elegans* remains to be determined. The lack of SID-2 seems to correlate with our observation that relatively high amounts of dsRNA are required to trigger the RNAi pathway by oral delivery.

The nearest non-target gene sequence within the *S. carpocapsae* genome represents an uncharacterised predicted gene (*L596_g5821.t1*). The *Sc-flp-21* dsRNA shared high levels of sequence similarity over a 21 bp stretch of *L596_g5821.t1* (20 of 21 bp shared), however qRT-PCR indicates that *L596_g5821.t1* had not been silenced, which could suggest: (i) the level of sequence similarity was either insufficient for gene knockdown; (ii) dsRNA was not diced in the correct register to produce this exact 21 bp sequence within a significant population of siRNAs; or (iii) the *L596_g5821.t1* gene is not expressed in cells / tissue which is sufficiently susceptible to dsRNA delivered under the conditions tested. In order to trigger significant knockdown of *Sc-flp-21*, 48h continuous exposure to dsRNA was required in the presence of 50 mM serotonin. Reducing dsRNA exposure time lead to a corresponding reduction in *Sc-flp-21* knockdown, as did a reduction of dsRNA amount from 5mg/ml to 1mg/ml over a 48h time-course. Phenotypes which developed following 48h dsRNA exposure were not observed across any of the experimental variations which resulted in decreased gene knockdown (shorter exposure timeframes / lower dsRNA amounts). This has potentially important implications for RNAi experimental design in other parasitic nematodes, and notably in *C. elegans*, for which the validation of gene knockdown by qRT-PCR is not common across the literature. Undoubtedly false negative determinations of gene function will be a problem in this context. Our data demonstrate that statistically significant gene knockdown levels are not necessarily sufficient to reveal gene function; careful consideration should be given to the design of RNAi experiments as a result.

The neuronal RNAi sensitivity of *S*. *carpocapsae* IJs, and the ease of behavioural assays makes these species ideal models for studying the neurobiology of sensory perception and host-finding behaviours. Within the Steinernematid EPNs, a number of species also display a highly specialised jumping behaviour which can be triggered in nictating IJs on exposure to host Insect volatiles (18). Silencing *Sc-flp-21* triggers pleiotropic effects on host-finding, lateral dispersal, hyperactive nictation and jumping phenotypes. The waxworm and mealworm headspace SPME GC-MS profiles are expanded relative to those presented by Hallem et al. (36), and likely reflects the increased sensitivity of analysis in this study. These data could provide a valuable tool for comparative analysis of neurobiology and host-finding behaviours across EPN species.

Collectively, these data provide the first mechanistic insight to a behaviour which may have implications for biocontrol efficacy. Through isolating genes and signalling pathways which coordinate these behaviours, efforts to identify molecular markers of desired behaviours and traits could facilitate the identification of more suitable isolates and strains for biocontrol use, and the enhancement of current strains through selective breeding / mutagenic approaches. The selection or manipulation of behavioural tendencies could lead to strains which are capable of operating within new ecological niches, expanding their utility.

## Acknowledgements

Waxworms infected with *S*. *carpocapsae* (ALL) were kindly provided by Ali Mortazavi and Marissa Macchietto, University of California, Irvine. Morris was supported by a PhD studentship from the Business Alliance Fund, Queen’s University Belfast; Wilson was supported by a PhD studentship from the EUPHRESCO Plant Health Fellowship Scheme; Warnock and Cox were supported by a Bill and Melinda Gates Foundation grant. Sturrock was supported by a PhD studentship from the Department of Education and Learning.

Dalzell was supported by a Leverhulme early career fellowship.

## Supplemental Data

**Figure S1.**
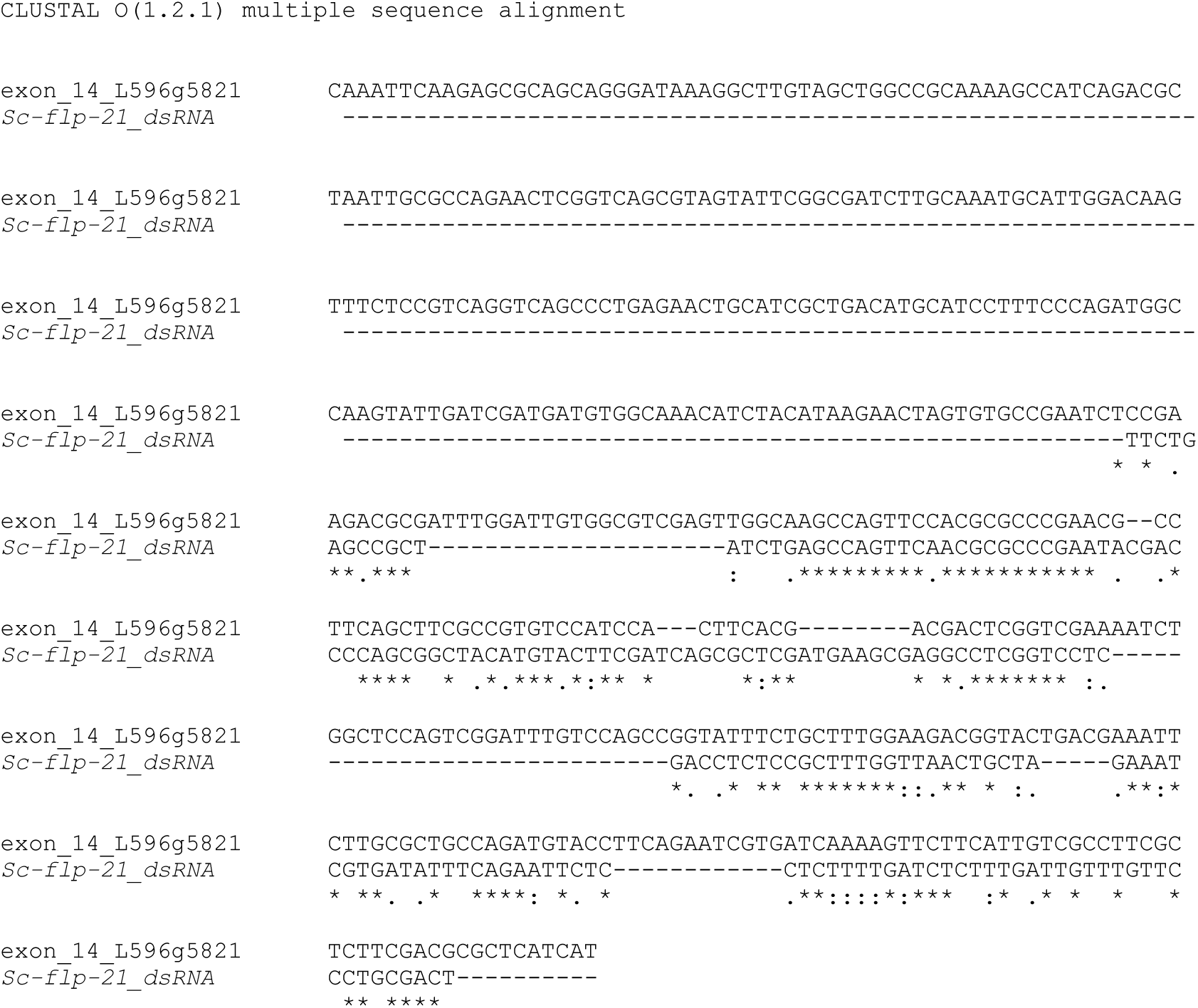
Alignment of *Sc-flp-21* dsRNA against most similar non-target *S*. *carpocapsae* gene (McWilliam et al., 2013).

**Table S2.** Steinernema carpocapsae argonaute proteins.

| <u>AGOs Protein List</u> | <u>Homologues</u> | <u>Accession</u> * |
| --- | --- | --- |
| ALG-1 | x2 | L596_g7718.t1 |
| ALG-2 | x1 | L596_g16709.t1 |
| ALG-3 (T22B3.2) | x2 | L596_g3457.t1 |
| ALG-4 |  |  |
| CSR-1 | x1 | L596_g20174.t1 |
| C04F12.1 | x1 | L596_g11107.t1 |
| C14B1.7a |  |  |
| ERGO-1 |  |  |
| HPO-24 |  |  |
| HRDE-1 (WAGO-9) | x1 | L596_g11197.t1 |
| NRDE-3 (WAGO-12) |  |  |
| PPW-1 (WAGO-7) |  |  |
| PPW-2 (WAGO-3) |  |  |
| PRG-1 | x1 | L596_g25491.t1 |
| RDE-1 |  |  |
| SAGO-2 (WAGO-6) |  |  |
| SAGO-1 (WAGO-8) |  |  |
| T23B3.2 | x1 | L596_g21112.t1 |
| WAGO-1 (R06C7.1) | x19 | L596_g19943.t1 |
| WAGO-10 (T22H9.3) | x2 | L596_g19923.t1 |
| WAGO-11 (Y49F6A.1) | x3 | L596_g17524.t1 |
| WAGO-2 (F55A12.1) | x1 | L596_g16917.t1 |
| WAGO-4 (F58G1.1) |  |  |
| WAGO-5 (ZK1248.7) | x4 | L596_g12936.t1 |

\* Accession numbers presented are the top returning hits for each RNAi pathway protein.

**Table S3.**
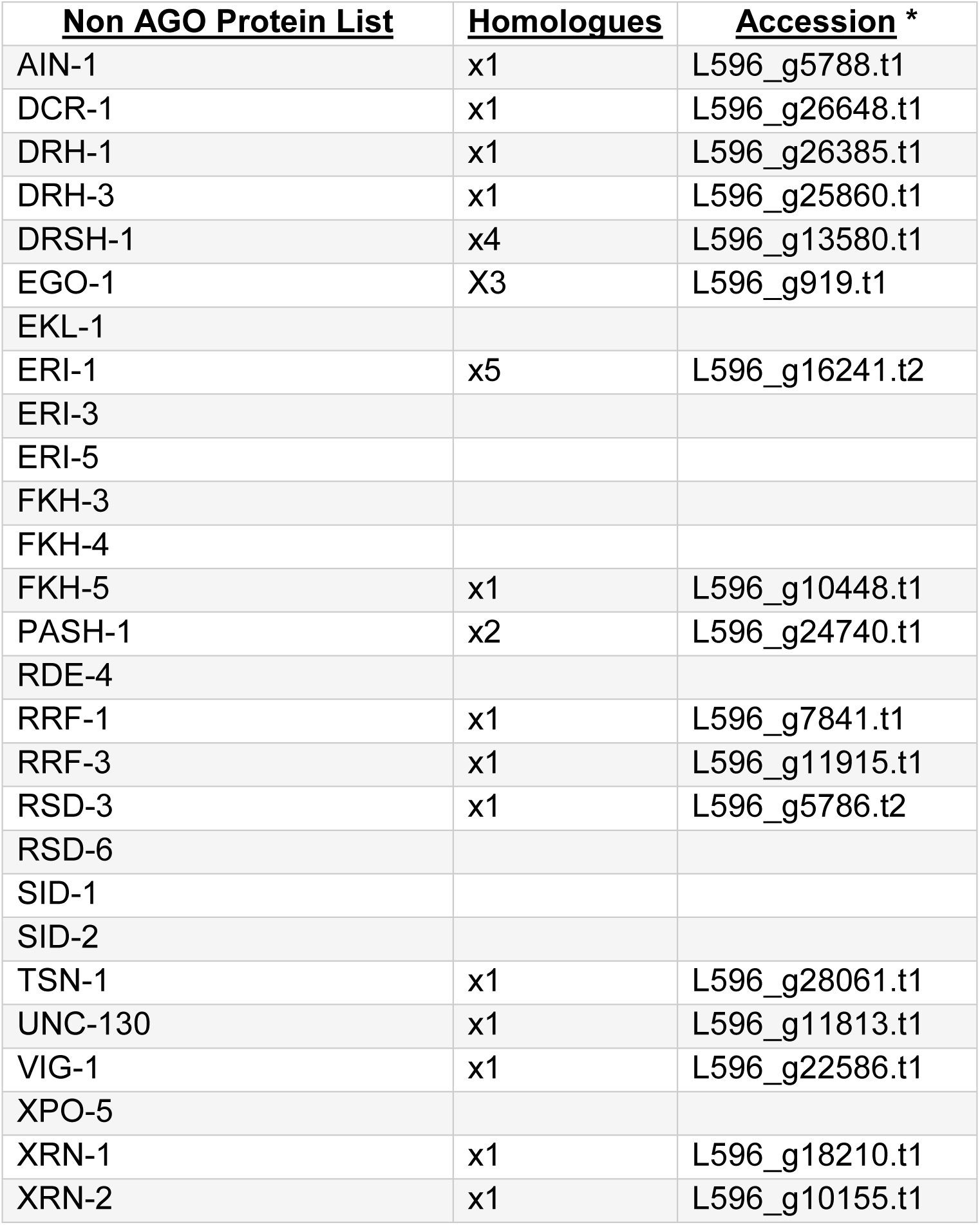
Steinernema carpocapsae non-argonaute RNA interference pathway proteins.

## RNAi pathway proteins

~~~
>AIN-1 - L596_g5788.t1
MWGEETGAETMTLNPWAGVSQQPQVWQNMNSRGNGNPSVANSVGSFAPNGGSVRPGSQQWDGGATPAGRSAWNPNPHAPRWTPSVNHGNASGWPNAQQHWSVANSNGTYAEQTKKNMALGRSSGPNLNMSQRFDGSQGWGSKVDQGTPWDTGNSGGVSQQMHEGKEQWSNPTRWTQPPGTWPTPIVAPMVADWQQGQPRHDHWANHPTGGMVPSGGAWAQQALGVASSSEQGRVPYDPNPQTPGPWVAPQPAEMNNDMMWHDPNPKQKKIQKDVGTGIWGDPTSQIEIRRWKDLEAEGGEFPGGSDWGSGSNSSTQPTGWGDVGPAQGNDGTDRWGAQGQALQGGWSDKVDQDRDLNGGKSDLNQIQRGIQNAIANGALCPDGQSRLQQWKKQRGGIGVGSPEEKMGVASSSDSNVPSLIAATKIMNIHGDWTPPSSVADDSAKSEDQVSSSGTNNEKESSVKESASPTPPPTQLDDGPQEFVPGKKWEWRDPNKVAEDPNATPGNCKPNPLMAAGNNMNFAFSNSAVNTANPYGPEVPSGNASTFWPNNSQGFNVPYGRDMYNSVRARLPSNGQFMPQRGMSGGYSNSNHNRMQTPPGKGVFIVLNHQGANETQLNFSCTRAGQLLNVASLGGQTVLLRYADSSSEGVLQKLKADFPGNERIKTVSEEEVEKLLKNRSPTMSGSAWAPNSDTPLWSNGGSPLATGEIMSSQNVFSNQEDLNRQQY
>DCR-1 - L596_g26648.t1
MSKFAGKPPQAPPRPQKLCGLKSTTRRRGLHFDGRAPAAKSRFQISLANFKFVWRRSGDSLADRLQKRDENWILKLRSPSDSAKAKNAAQMVVKATEINPNFFTPRDYQIELLEKALNYNTIIPLGTGSGKTFIAVLLIKEYTQLLLPKFSNGGKRAVFIVDKVALVKQQADHIECHTELKVAQFHGYMNTDVCNTREAFEKVIEDSQVLVLTAQIFLMLLDHAIFSFDRVPVIVMDECHHVLGGKHPYRLIVQRYSELSEETRPRMLGLTASLINNKTAPSELEALLRQLETIMKCRIEASSDIISVAKYGARPKELVIACRDNDFAEDWIRVCLKSLQNLRESAANCLDFHPDLDVDPRKSVVEAATKILSVLQQMGPWCAWKVAMVWEKQLRKIHTSKSKASGLGEKQIAFLQNGESKIHEIIKEFEQKVKTVRSYEELKKYIPHKVERLLELLRYYHPNIQKTLQGGGVKSLSAMVFVDQRYVAYAMNLLMKSLYKWNQIDFGHLRSDFVIGFNGTSLGEESAGFHKRQETVLEKFRQSQLNLLFTTSVLEEGVDVKHCNLVIKFDSPGDFRSYIQSRGRARKQGAHYFMLTEAKNKLEFALNLRNFCEIERMLITRTVTIHDPEHQVPARNVDDVVQPYVVTSTGAKVTMSTAIALINKYCAKLPSDVFTRLVPQSCIEPVTVNGVTKFLATLSLPINSPFKEEIRLKTPMDTKKLAQMAVALEACRQLHEKNELNDYLLPAGKDAIAATFLLDEDPDEYVPHMPHKAGSIRRKQLYDKKMDLLVEASGAANPKRRKIANPLDNDYFFGFLSKRILPEVPSFPIFPRQGKVLVSIKLCKNQITLDEETFERAMKFHEYLFDDVLGLAKQGFVEFVPRHAPIATVIVPLKRREHAVYDLDRVYLSNILEYRRIPYTPSEEERRTFQFREEDFTDAIVVPWYRNQDVFYDVAVINHSLTPASGFPDGNYTNFREYYMKRYNLEIFNETQPLLDVDYTSTRMNLVMPRIPTRSQGKEKKSDRTQHQVMVPELVNVHPIPASLWNSIVTLPTIIYRANALLLADELRELICAEALGPNDNSVHICWEPLDYVTSYSEDAQLPINKLSHLEKDLEAERAKARAENPEPEIVDMEDDSSNDGFNIGVWDPKLGDGGELAGRILTVPFPMASSARTDEEIADEAADMIVTSGNLNNGNMSDDEDDQGEMKMLMDYAHIYKNDSSSFLPVRDNIEELGWDADIGQLNISEDAFPVSITGTSASIDGKSLMKDIASVLDPSFAGNGVASANGSAPNGSGDAVFAKSKPVATRLDLSAFDDRDAPLIRTGENESLNLIEYFMDSNDSQEVERMKKASLEIESSRPHLENFPVVEVEEEGTPQKAIEIDNSLFTNPNVLQPEINAYTDGDVGERSEVARVIPKTTFMKPDVKVNFTDEGMPECSAGVSPCVLLQALTLSHASDGINLERLETVGDSFLKFAVTDYLFHEHKEQHEGKLSFARSKEVSNYNLYRIGKKKNLPSILVASKFEPTDSWLPPCYMPTGDFKGPNAEDAEATDKFMDAVFEGKPIPQSKKPLTGWDEDNNEAEKVVDGIETINLLKNPTGNTQYDVNEEISPLPYNVMTQQYISDKAIADTVEALIGAHLLQLGPKVALRFMRWLGLRVISEQTVADEPLLRFLDTPEDPNRSNQELVSYIERFQLSSVEKTIGYRFNNKAYLLQAFTHASYYKNRITSCYQRLEFLGDAVLDYMITRFLFQHRKKYSPGVLTDLRSALVNNTIFASLAVKYNFHKHFVAMCPGLHHMIEKFVRLCESQAKNTNFNSELYMVTEEEFDDGDEEDIEVPKALGDIFESLAGAIYLDSGRRLDVVWEVFYNLMHDTIMECCEHPPKSPIRELLEREPDKARFSKLERIRENGKIRVTVDIQGKCRFTGIGRSYRIAKCTAAKRALRYLRDLDKEREAAKKASAHH
>DRH-1 - L596_g26385.t1
MNTHLKTSHPLSVINWDKFTPETPVAIRPEHFDLRPYQEELAHHALKGENTIIWAPTGSGKTIVAVHVIAQHLLKGNGRKVCFVVPNVTLLEQQKRVCERHMDAKVNIIKAESKAPFSGMVAASHIVLLTPQMLVNALQNTTEGKENFSLSVFSMIVLDECHHTAESHPYNVLMHHYHDVKLKGTGKSPQIVGLTASLGAGTSRNANEAFRHIQKLCANLDSPKISTVEKHKADLTGFAVETKDRIRYVESNYRSKPFFKDVMELMSELENGIFVHPAIACDPVKKAMIKEQVCRSSDASSTANYRSSASTRYSQAYQNWLGNALMITLPKVDLPSDARVEIMTRFRLLKILYKMIELWANFNGKCAMDYYMKESEGLPIPLPVHDRIRELVSYRNDNSDMLIEMCQLLVEKFGEGTDGASPARVLIFVKEREYTYMLANIIDCCNELKALGVRPDFLISTNSGDGECRLNPEEQRSKLEKFRSGEINLLCSTSVAEEGLDVAECNLVIKYNYASSDIAHVQRRGRRARHQNSYAVLFTHDKNLEQQENKNILSEELSNQAIALLRQMPQKMFEAEVQELVRKSYSDRVIERAAKETASLALKDVGARTCTEFIGRSTDVRHSNHSLHILCDGSIWDRFTCEESVSQENFLKTSELPIGKIYCNNCKEVWGRIVIFKGIAVPIIACKGILLQNARGERFPCAKWAGIKGRYFCPEEVKSVDLARLSAAPHRPQFLTVLMGETESRSL
>DRH-3 - L596_g25860.t1
MTSNGTRFELDYVLTVITTLDFILEPFFLFLALTKSTPSMWFYRIFLIAISLCNCAFSIVFFLLSAKFIPIDDAVCLISSHFALRSGRVLWQLSMFFMMIQFQIVLLMLIYSSYEISHPLRPLNWRSKAVILAVFVFIFLPASLVLILEQQVIMGEIEEEKREKVPLSHVRLFRKEFLESGRINEAFIDKCEKFLSFDVVEILKKLFTINPGLARKFFFRQIRLKHPKESGIIIPIIERCYGDDTEVLSELRLENLKNASKERYLKIFSDDEFCAGDTLSTQIRPQPFIKNLAAKYGNRYEKMIKKVEALLKDKEHDAAACQILRSMPKIADDDEDDSWWLHLLDICDMDPHNRACLVLLDVDYRNIIDMFRAEQRAKRTPAIQGKDEVMFDETISDLDREFRGQEPYMEHRETVDENLIDDPDKIVLRDYQEEQMEGVRKGNTIVCAPTGSGKTIIAAQTLLEHFKENEKARAVMFVPTIPLVEQQAQLLTRFLRKRKYILTASGAERVVSLGRKMLSADVVVITPQLFYNYMTDPRESERIYISDFTLFIFDECHHCDSNHPYRKITKRISAFEGKKPHVVGLTASVGVGTKTMNPQEAVDHMIRICANMTAQSITTVQRCLKSLDEKVRIPDDYIRTVHRDLESPFRGKIAEVLGKFVYIAEEKLKNTVLAGGEFESFPDKRKVCQFFGFVKRLKNELERNQKVEDRNALIYTLARIEVLYKLFNLSDLLPARYAAHVLKEFNDENKDGLQEKYKHYLRELGEAAIKNEERDMNKDILVTLKEILVDQYQKDPFSKTLIFVSTRDLARDLSLYLRDQWDKYGLPRKVKPTREGEDSEKTYKPVSYITSSNQATSAGGLTAQQQRETVAHFRSGHHLVLVATSVADEGLDIAECNLIIKYNTSESSELKLIQRRGRARAKQSSSILLALDGAIEKAEYDGMLREKLMYAAIKHLHTLPDTHVYRLIEAKAQQLLVDEKEAEERAKQRKEEASEEVWIVACKKCSEQLTTTGSLRNAGDKRVASCDPSIWEKLKIVGNQKAESHGNASLEISVAVCSNGDCDNTIATLGTDGNTFMPFLRAQSVSLYRQAEYDADNREGKQSLKKWRSLEAGEYNIVLRSIDDEDRKRMCSSWESQNSEKAKEIVQKRQDQRAFKRARYLTKESRNRELAKKVILKQKEDPNAEEDDYGYSSIADDKIRGEDLEKVKIKAEDFAQVLHSEPNYLSAMKGYEAEVSDNSDNDYE
>DRSH-1 #1 - L596_g13580.t1
MEDPGCSGTQKLFEGAKRTLDKREDAQENEKTENKEQLLTESDVFKRLAEIDLTTEESTANSPDPSKIQDLSKFTVAVSKRTISEPDASEFYVKTENGDTIGTDRLKTVHEAMQFRVLDVIDMARREQPQSEPPTMPTFHENCKCGHVSESEADSSSGSESEEDDENADLSNVSDANKHSLVAKREIARKQQHPAGLSQEMCFNEPGQMNDGPACRCSWASKQSGIRHSIFVGEERVEPCNPMSSNLRNLHHYFMKVTPNPVEQSRNASLIYHNDKGYVFEGFSVFFHRKLPQFFPQTPVNKLSQEFEVVFIEEKAPEQFTVHDLESFHTYMFDHLLEMYDLNRRAKDVFDGCPVYHCMPRFVFKDYATEAVEVLPMSAVLHHIAGAFQPVFDDRLVQKIRRDENLFHNMSSHMKGQIFVNPSKRPATIRVDLIERQDPYTKSVNKDFHPVLTHIGLRPSVYTFSAKPAYQQAMKKYLRTRHLMSLQGKLSYEDKQKLVEQEANIRKLKAEAQHKRDMVMSVTSRGFYSTTFYPDIVQHAVLLTLACSHVRYHWCLETFEKRIGYSFKNRTLLELALTHPSFRANYGTNSDHTRNALANCGLRIDKARNDNRNSQVDRPSRKRGYENLREVMSMKGTEKAVLSPVHHNERLEFLGDSVIEFITTIHLFYMLTDLDEGALATYRSALVQNKHLAVLAKKIGLDEFMLYSHGPDLCHESDFRHAMANTYEAMMAAVYLDCDLNECDRIFADTLFMDEKEEKSKEKLAWTKLLDHPLKRDNPYGDRHLIPKIDSLQLLTQFEDSIGIKFKHIRVLAKAFTRRCIGYNNLTHGHNQRLEFLGDTVLQLVTTEYLYKHFPNHHEGHLSLLRTCLVCNRTQGVICDDLAMAKYLVIPPNSRKHTHVMSIRWKERADLVESFIGALYVDRGLEYCKTFCKVCFFPRLKYFIESQRWNDPKSQLQQNCIALRDGKNEPEIPEYRVIAIEGPTNTRLYRVGVYFRAVRLADGVGHTVHLAQMNAAENALKQHADMWPSMSTKKTEKKASHNNWERASYRREGNSAKYERSSNRQGGGNDYRNDLSGQDYRKVPDGRESRRNSQNSDAPRSTFNAPYMQQNRRPYDNGQEKPHRSRNGQQYDDRSNQNDLRNGRNHHAPSYQNEPSRFGPTRHRHDHEFRKPYDRPQHDFRHFTNDRRPHNESQNTSYGRDQNKGEAYSGVRSRYYESQSERPPYRPSGSQKDSNPVYQRNAQPYRQQNNEQNSCRQPTWQDSQNKPH
>DRSH-1 #2 - L596_g13580.t2
MPRFVFKDYATEAVEVLPMSAVLHHIAGAFQPVFDDRLVQKIRRDENLFHNMSSHMKGQIFVNPSKRPATIRVDLIERQDPYTKSVNKDFHPVLTHIGLRPSVYTFSAKPAYQQAMKKYLRTRHLMSLQGKLSYEDKQKLVEQEANIRKLKAEAQHKRDMVMSVTSRGFYSTTFYPDIVQHAVLLTLACSHVRYHWCLETFEKRIGYSFKNRTLLELALTHPSFRANYGTNSDHTRNALANCGLRIDKARNDNRNSQVDRPSRKRGYENLREVMSMKGTEKAVLSPVHHNERLEFLGDSVIEFITTIHLFYMLTDLDEGALATYRSALVQNKHLAVLAKKIGLDEFMLYSHGPDLCHESDFRHAMANTYEAMMAAVYLDCDLNECDRIFADTLFMDEKEEKSKEKLAWTKLLDHPLKRDNPYGDRHLIPKIDSLQLLTQFEDSIGIKFKHIRVLAKAFTRRCIGYNNLTHGHNQRLEFLGDTVLQLVTTEYLYKHFPNHHEGHLSLLRTCLVCNRTQGVICDDLAMAKYLVIPPNSRKHTHVMSIRWKERADLVESFIGALYVDRGLEYCKTFCKVCFFPRLKYFIESQRWNDPKSQLQQNCIALRDGKNEPEIPEYRVIAIEGPTNTRLYRVGVYFRAVRLADGVGHTVHLAQMNAAENALKQHADMWPSMSTKKTEKKASHNNWERASYRREGNSAKYERSSNRQGGGNDYRNDLSGQDYRKVPDGRESRRNSQNSDAPRSTFNAPYMQQNRRPYDNGQEKPHRSRNGQQYDDRSNQNDLRNGRNHHAPSYQNEPSRFGPTRHRHDHEFRKPYDRPQHDFRHFTNDRRPHNESQNTSYGRDQNKGEAYSGVRSRYYESQSERPPYRPSGSQKDSNPVYQRNAQPYRQQNNEQNSCRQPTWQDSQNKPH
>DRSH-1 #3 - L596_g28273.t1
MTTSPDIELFNISSDEEDDLLDFPKMAPARKATQAPSREKHCQAELSRACEAFSVQTLDAIGVARCRAADGNAAENGFETLLGCPCPKSQLGTGLRHGFYPKEECIPACDKEQINPPNLHHYVLKVDSPNVPQVTRIMHGNKEFVFDGFSLFFHQKLPVELLCTPLKIALGTSATVTLLPARSPKGFNVKSLDWFHKYLFEEILSFPMSQPNQNQCSFYHCLPHFVHKEGKNVHLLPMIAVLQHLQANFKPIFGVNDLFKNHRGENVVAKIRGQVVFNPTKTFSALRLDHVEDPRDLLGQPKLVSFTKWKKSVHTRSRTKRKFMPNRIKSDQTERPFFENAVPSKGYYKTGLYPDVIQHALLLVPTVNYMRFHFNLQILEVRIGYTFKDKTLLELALTHGNFRGSYGPHRLIVKRQNETFGFRTNLDTATKELKRPSSNGSKREPGQTYERLEFLGDAVLGFIAATHLFYMFSMADEGTLSLFKSKNVDNSRLCDLAKRSGLQQFILYECDRNLIIDEGSHPLLANIFEALFGAIYLDGGLNECDRVFADITFRNEQNGEENKLAWKNMLEHPLKRENPHGDRHLIPKCKSLLLLTKFEKSINAKFKHIRVLAKAFSRACIGLNNLTHGSNQRLEFFGDTILQFLTTEFLFHKFPSHSEGALSLLRQGLVNNKALSTICDKLSMRTYIVVPEDMEKMEHVWILTEKSKADLVECMKRWKRKAGRTRNPPSKKRSTSSAEALDAQIYLDTKFSATRDHQIIDLTMSPSTSGTFNTLKLFKSLFRGKQLGVGKERTIKKGEMEAAKNALESITEAVFKDMKRKLLEEKKIDVPRTGVCSYPSL
>DRSH-1 #4 -L596_g13580.t3
MEDPGCSGTQKLFEGAKRTLDKREDAQENEKTENKEQLLTESDVFKRLAEIDLTTEESTANSPDPSKIQDLSKFTVAVSKRTISEPDASEFYVKTENGDTIGTDRLKTVHEAMQFRVLDVIDMARREQPQSEPPTMPTFHENCKCGHVSESEADSSSGSESEEDDENADLSNVSDANKHSLVAKREIARKQQHPAGLSQEMCFNEPGQMNDGPACRCSWASKQSGIRHSIFVGEERVEPCNPMSSNLRNLHHYFMKVTPNPVEQSRNASLIYHNDKGYVFEGFSVFFHRKLPQFFPQTPVNKLSQEFEVVFIEEKAPEQFTVHDLESFHTYMFDHLLEMYDLNRRAKDVFDGCPVYHCMPRFVFKDYATEAVEVXXXXPKLQLLLLVTVCAFH
>EGO-1 #1 - L596_g919.t1
MLAAENDRPPTSDDSQMELKEGGTAVMDVKARDAPSKKAPRVFPESDDSDLVRIITEAERDRRMQLKLDKADFQKKVIHEMSADIAGCSASCKRVSEVPGNVNGVPKSAHFEVELTAPEWDRAFFPMVHNFLQISKDYAETKKANALMQVSNPTLLREDFEPTNKLLPLQSFAFGCMRSPFEFTNHHEFHSSESNAELLHDYFNHCREEPNHKEAMTIDFNHDLGDSFVKFASEEGNGYFVYMLKLIYNKVRRIIVNFERLDSTTSTVHGVKLFLLFNNPIEVRRSQKIKGRKGDSYWRFDTNGDRCLTWNNNEERQAIIADCLNLKLQFEKLEKNVLYNILSRLRRRCNVILEFSSVQEVPFKGHVDRPMFGDHAIARKLKEMDKYDVNYLIEALCSRGAVINDQFLESEKTRDDYVEMILEDYERNSFITIAALERILNIIDEHLEVRDIHLLYRTCFNEEAAHEKDLKETEERRKADGFLKVRKIVITPTRVLFVVPEMMMGNKFLREYDPDGLYSLRVAFRDDSGLKMNNSNVGSQLVEKTIRESFKGVMVAGKEFVWIGSSNSQMRDHGCYFLLNQTRDMRKILMEKTGEFDEKDPIKAQARFGQRFTQARAVPFDLSRDDYDFFDDIKGGEDENGERYTFSDGVGSISVHFARRIAEALKLQNFVPSVYQIRFRGIKGIVTLDPNLDYYRDHAKKYGIQERRKHHRFDLQIAFRKSQDKFNARVDVEIDIVKFSTPTPMQLNRPMINIMDQVTAMQSEASHRRMCARVHQLLDQQLSDLGKCLVDEHKAREKLFDIPRRVDVHLLTLEKGFHLTEEPFFRSLIQCHVHTTLNRLRAKNQVQIPFNKGRMMFGVIDESGGLQYGQIFCQVTNNLQLKTPGVTAAKTILTGPVLMTKNPQNVAGDMRMFNAVDIPELRHLVDVVVFPRYGPRPHPDQMAGSDLDGDEYAVIWDPELFFDRNEEPIEFPKPKKPVPPADADSTDLVINHIVNYIQQDSIGVISYAFLVNSDLYGIDSEVCKRIARKHSMAVDFPKTGVPPQKLTREPDEDNNVPPEKPDRYPDFMEKSGRDPDYVSAGLNGQIYRRSKEIQGILQFTVGHQESRAPEPDPDFLVEGWKTYELDAREAMDAYNASIRALLDNYGVKDEAELFTGNISNARNRLSDKDNDDMSLFNTNYAIEQRVSTIFQTHREQFFDKCGGYMELTVSDMTFIPTAPKINSRDLERRLCTEISPEMKKRASAYYQVCYTLQPKRESRRILSFPWIAYDVLAEIKKENYARKIRNQELRIDIGRPIMSRLDDYIKEYLKHPRFAAFAKALVFDKVCHRYATVYPALTELFFVLWQFGFEQKIFDGCILPENLFYLFLLFAVSTENPFLEKIPADANVKAMEKRDLHARIGGMGGPLMDFLEYLCTREFQSQLFVSFNEIGVDDCLQGATLASLSRAAIHTYYRITFSGMFFALPQLGADHSKNPSTKEQKFDTSGVRVYEIDPFVIEVPSFAKEEGKLDELTQMLKLRTGMQHLVVRIKFIVRKTGNLRLLVSGIGSLEAIEALRDVVSIKPNLRSGSATVVAKAHLMADMVYAKIMGSDNFGGGPFGYCHDISDIKKATPRKKPIRKDLRN
>EGO-1 #2 - L596_g10626.t1
MSADSDSSANAKTSGVIRIFLDRTYERYDRCTELQHYVLEKLTREETFQTFTLKRNGLPRHHKGDYEERLSEYELTRMPPPKNADKSKEPWDREEYPPRKRLLQLVTEFTEALSATMKGPLPAIMQISDDFFLKKDYDPIDENLRVAEFALGNLVDPFTFHYSELKSSTDDKAQHDTHILRSFMKDNNIKSGMIATFEHDKNILWIRCVHWQKNPQAVYGVSLKVPYNQIRRLIIDVDKKADCATIYFCLNYPVEIRRAKMKSQKGRNMSAFGTEFDRKGRDGERCLSWEGEDSENGRLKQAIADSPVLALTLTDYKDEGFYNVLSRLRSRCKVPEFTHVHKEKRSDYVDPPDVLLHLRTKSGKLNDFSVAYFVQAVFSRGALVKDALLRSEDVRNKFIAKICKQCDKNRIVAVCALERFLNAIDEKRELRDPIKLWDFLVNQVSGDIGTIQKQLDKNEKDGYTLVRKAVVTPSRILLLTPELIMGNRGLRDFVNNSDDVIRVLFRDEDGTQMRKISVGEIIIQKVVGQVLEDGIRIAGRHLCYLGSSNSQLRDNGCYFFEKSQIPDIRKKMGTFKEMSVPKRMSRMGQFFTQAQRVQRPLERKEYRESYDLIGGADLVNEPYTFSDGVGVMSVALGEEFSRSLKLNGCVPSCFQIRYRGFKGVLSVNPMLDEIRAWAEANLSDEDIPKYRNNVVFRSSQKKFDASKNAKLDSAPFEVVKYSAPCQISLNRPMIDILDQVSEIQSYKAHERIWSRIDELFEKQVAKAASSLTSEHYARERLAELPKRIGTSDLKEEDGFMLTQEPFFRSLIMSSVRLSMSKLRKKNNVDIPSNLGRLVYGIVDETGQLQYNQIFCQVSSSIFVKHPKKSALKRILKGPVMMTKNPAIVKGDIRRFEAVDIPELRHLVDVVVFPRYGPRSMPDEMAGSDLDGDEYVIIWDPKFLFDKNEAAMNYPRPLKTFEKVREDEIEKKSCEFFIKYIIQDAVGMLAHAFLANSDYYGIESEVCENVARKHAKALDFPKTGDQPDALTKMPTKSEIYEGKQIPQEKPDRYPDFMEKDHQRCYASIGLNGHIYRRARALDDILHRSVDRHLSDKIEVDPDLIVDGWEDYEDEAKKTMNDYNSQIRSVLETYGIEDEGQLFSGAISKMRSGVGDATQDKDLFGAFNVLSTVSKHVALIFDEFRRSFFDEHGFGGYLHCTDLKYETVGRSDEMHHSYKRTCRKPTLEMKKLASAYYKLTYEAARQESQQAVKLLSFPWTVWDVLAAIKHDVAEEKRRSSDRRDQYMNRRVKPFANLLSEHIAKFCRQRQEDLDDFKNLLVNTDGIIALMLERYEGLVSILFFVCVWADDSNVFSDFFNYQHLCWILILFGQGIYPWNHAKTFHTDLPIFYPIDKLKEVQNIGKESVNLNHLLGGIGSVILLFLEFLSSYYFGHKMKSISFNPLGLNSSLGSCYLQKLFHAATKTYNEVIFSVSFDTLPQPEEGKVHRSLREFEIEPFMIELPLQKESEGSEARYNLSDVEDQIKEMTGASHVRVRPLNDVFLSGVTRLMVSAKGTLESSQKLRTLLSVMPSCRNEHNAKWGKLFMSERLLMRLNLVHS
>EGO-1 #3 - L596_g1037.t1
MHEILFLLAEGFVEGPVGALEGKDLTVFFSKSFALNFSWIKSPPVTINRLTLCVHQIDSMLNDETKALARLSELTRRIDFNHLTSEKGFQFIDEPFFKSLIQCCVKFTLSKPMGRFKW
>ERI-1 #1 - L596_g16241.t2
MRDRLKRRLRKDLLSTREYEDEFSRNKLRQHFDFLVIIDFECTCEDEVVDYPFEIIEFPAVLVDTHQKKIVSRFRTFVRPKVNPVLSAFCVELTGVTQADVDKAPRFSEAVLKFHRWLFSHVYPSGQHDFGGLMKSYAIVTDGPWDLGKFFQLECIRLECIRSERYNAKIPHHFRAYINIRRVFTLKYKKTHALSKVNLAGMLAVLGMEFEGREHCGLDDAMNLARIAIRMMEDGAEFRINEKLVAAEYADKYTSFVPTISTTGMSTAERDRRKWRLNLPYRIVNICKDRFMTGAYADCDSCDEDANFYEKAANRK
>ERI-1 #2 - L596_g986.t1
PISAHNFRGTSQQMRDRLKRRLRKDLLSTREYEDEFSRNKLRQHFDFLVIIDFECTCEDEVVDYPFEIIEFPAVLVDTHQKKIVSRFRTFVRPKVNPVLSAFCVELTGVTQADVDKAPRFSEAVLKFHRWLFSHVYPSGQHDFGGLMKSYAIVTDGPWD
>ERI-1 #3 - L596_g8761.t1
MSLEELSALISEASKRKPKRAKRGEEEAPNGRYEDPKEFPAVLVDTHQKKIVSRFRTFVRPKVNPVLSAFYVELTGVTQADVDKAPRFSEAVLKFHRWLFSHVYPSGQHDFGGLMKSYAIDDDGLWELRNFLQXXXXSILTGLSVKKRNSAFIVQVRTTYPMHTASTDGLAKMSWGSDEGSISMAIAALRKSRPAAFELEYPDLMAKTKPTPLPRQPSPAQQVEPKSPAQSIAPTSSSEDLHAHLAKIQVTQPSVYLPSRPAYPSVAPSVPTPEAVVPQPEPPRGPPGFDRSAKPKETTFDTRVMGGASQTDVQVVPSREVPHVPSLPVAPARPRVDRSTKEAVARKEAEFQRNLFAVYEACTQDFRNMT
>ERI-1 #4 - L596_g16232.t1
MLAVLGMEFEGREHCGLDDAMNLARIAIRMMEDGAEFRINEKLVAAEYADKYASFVPTISTTGMSTAERDRRKWRLNLPYXXXXVYCGNAIINGRLIRITFKFSRIAWSQHLACEGVTKIEMLAAYVAFEARRRLAQFVTYRES
>ERI-1 #5 - L596_g16263.t1
MSVSVTRIQELPSVLRPFVPRQSLPSFSRDGRTPETELRSGKTRGRLLRVRGRRRGRRRGFRRRRVRLQPGLRRRSDDLRGTLGAYLRSLETDAEESEKRGRGCSKWPLMIEDPNEPLRCEGYQVPKHDFEFPREVSVXXXXKANLQKELEELQLNTGGTSQQMRDRLKRRLRKDLLPKYKDEFSRNKLRQHFDFLVVIDFECTCEDEVVSRFRTFVRPRVNPVLSAFCVELTGVNKAPRFSEAVRDEFSPGRESDEGQGRVSDQREARGGRVRGQIHRVCPRNFDHRHERRGAGSQMMWRLNLPYRIVNICKHRFMSEAYADCDLYDEDANFYEKAANRK
>FKH-5 - L596_g10448.t1
MTPRGLCVCNLSSQPQHSMEPSSESSRKFMDMISTLCAEVDNKQKLPEKKPQRNASGKCMKPAYAFSAIIALALKNSPDGRLPVWKIYAFILEHFRYFRTANEAWRNSVRHSLSKNGNFEKCLKEDGSKMKDGNMWQIIESRREAIDREILKYRLSDKYHAMNLAALNKPEILTSLEEGSFGLPPFVFEKEYKQAYRILANQKNKRPEIPKNFEVASNVFKESNMNTLNTDLDTLARPNPISVLLNENTPSPISRIDLLKPSIPSSSSQNMSEASMNAEEKQIYEKYLTSDSNPLLSSAALPGAVVPEIDDCMLWECEQDTLEAGIQTRLRHDENDFWLPTRSNCIFDLPSDNLSVVSDGF
>PASH-1 #1 - L596_g24740.t1
MRKHKKLKPRILLAKTWCSPSPFGDLXXXXSCRKEKSLTTPTDHLIEDSDSDGGIRNRSFGETQDQPQTTQTDHLDGSRSGDEFEDETLEKTPNKSHTSKNEKEEQGTSKRVLAYSEYGLSNKLPEGWVEVSHESGVQVYLHRMTRVCTFSRPYLLGEGSVRHHKVPQTAIPCFFQRKVREQLEQREKEIEEVVKSQKGAETSAHLIAKLQAPDVKTIAQSQLTPDDLFDYAKSVFEFKTITVYRDKASKWRHLKEQKRAEGKINAAAKAALLDSAPEIPQNMKIITIALDVESEKKAVTIDPRAKTSINILSEYFQKVFRSTPRFTDELTSVPLKEKIMSFQNLAKSELNPEIGDEVILGEGVGAKKKTAKLRAAAEAASFLIPEIKFDLDGKVISWNEDSGNKTEKDITVDFNQYEITDPRIAELCKSYAQPSPYTVLQECLRRISACSNDEIQIKTKHKNYSSFIEFTMAVGEQKVSVTCSNQQEGTQRASQLMLALLYPDLQNWGDVLRLYGEEPQRVVKDRSKEVTTLQTFVQNGGDKVLSDHNNAILDKLKEEMKATADGLDLQKGLLKEVSASHLPISECDAVLSAVYAEFHRVKREVPEMGI
>PASH-1 #2 - L596_g24745.t1
MGDVSDDLKLLMLERERILRELNSQNPGYDPDDVEPPPPPPPELPDEESSLSSAEDMYPPPPPPFTESFPSPSCSPAAPVCLTASPANPEPCTSFDSVSRDPSSLQRDSSSTALPATPLQAPIKKDAEESFDSPVAKRPRMDGCPFSGTQFSEESPAVNDRKSTYNDDNVVSDADVSNEANNSQAQSVENSGDSDLSDDSDDDFVDGLLEKTLDESRTVKNEEDLEIKSKRVLEHRGCDHFDVLPEGWVEVSHESGLSVYLHRKLRVCTFSRPYFLGTGSVRHHKVPQTAIPCFFQRKVREQVEQREKEIDELIKSHKETESSAHLIAKLQAPEVKTVQDFEQSQLTPDDLHDYAKTVFKFKTINVHRDRTWASHRMRIKDKKRADAEMNAEANAASLDCDRPVLPSDVKLITIPSLDQNGKPQRKNFLMNPQGKTSVSVLHEYVQKVVKSTIRYHYEETRSSSNPYIAIAYLKLGQISGKATSAVSIKEKIMLLHEQQRREGLKNSSENTDEVFLGKGSGVSKKLAKLRAAVEAVSILIPGIEFDSEGMVVNSKKETGTENNANEDVISMFNKFDITHERIPELCSRSGQPAPFLVLQECLKRYSASADTKLSISSTRQKHHRHEFVMSVGKHETKVICTNKQEGKQRASQQMLQLLHPELKTWGEIIRMYGYEAQRQFKDARKKGNDVTKLQALAADGSSKVSLNPNAAILSRLNEEMRKTAERVGQLKGPFRPDLSNDETAVTDWEVALSAQYSEIHRDRVEVPIIDL
>RRF-1 - L596_g7841.t1
MDDMSMFNTNFVIEERFSKIFQKYRLEFFEEFGGFEACTVDESSLVNRSKNLHANKAESDMDRRICKNPTEAMKDKACAWYNVCYFYANKAKRRYLSFAWIVWDVLAEVKRENHFKNNREQRLMGIPIHTRLHAYIEKYTGDVSNRVALEELKKLIAKEERHIAKYVEAHEGEIWIVNCS
>RRF-3 - L596_g11915.t1
MSSKAVSTRGVAFALRLSVSQEGLPKEILLHAARQMLQSCGCSSVRLETPVQALQAEYEDCRLEISGTVTFAVPSSVDAWQMIIVFTSKFCAETGMGLELQPSLEMDAFDRPNHDNCHILWFAFGNMPNEGLFLTRGDYISGYNKKSNRFVPDRGYNVNYVAGTQLLLSWANFEHDRKLLTIYFAVQLPCPASDGLLFKGYKLVFTYHNIISVIADTDDSRAGNNVVYLKLRHPPQLWEAIPRLYANRRLVNLEACRDWIRVFEFPGSNRFYGCTKSTLGSSSVFAFGMPKNVVDPKILFEEEREEWKSFAEDLTTRENPTRSLYDILSRLKRKANIRLYFGSILSVVRSVMRTCDLPSTDSFRVNYCLEALASRGFSVMDQWFPIDNQEANYFPVFFSRVVWCLGECKEAVENTLENMLSIFDERKHHVNMVTVFEYLYEQNIKSLVEERDMDDCSYNDLPTNCVMVRKIMVMPSRTLLMPPEVMMTNRVIRQFGEENALRCVFRDDGGNKLVPKEFTRGRSVEGQSVTIKEIVKGTLSSGIVISDRHYRFLAWSNSQMRDHGCYMYADIITQDETTGEEIVQDITTIRKWMGDFSSSRNVPKLMSRMGQCFTQAQPTIRLGPSHWCVEKDFFTGPFGNPTKYCFSDGVGRISLRYAERISNLMGLTFSPSCFQVRYRGFKGVLCVDPQLDRTSDTPIVFRESQMKFIDYERGSQGPVLEVVKYSMPSPVCLNRPLIMILDQVAEKNGSHCHRRVCSAIHQTLENELNELAVMLYDESAAIRALSQRVNLSIDFGQFVNTGFRLTEEPFLRSLLLAIHKYNIRQQLSKVKIPLPNGMGRTMYGVLDEYGVLQYGQVFIQYSQSINRAGQRKILHEGPVMVTKNPCHVAGDVRMFEAVYQPCLEHLVDVIVFPRYGPRPHPDEMAGSDLDGDEYTVIFDTDLFFGSNEPAMEFPKSEAPEFDVMPETDDMVDFFLKYLEQDSIGRMSNAHLMMSDKLGLFHEICNNIARKCSVAVDFPKSGQPAEPLQMDEQCADCPDYMKSNTKPSYRSKRLIGQLYRKAKNIEDIIDLMPNPGTDRNIPFDEDLNAEAYLEKNPDVLRECIRVRNAYNCKMQQLLDEYAISDEASLITGHIISMKRLTEMEKDDYTFYHTDRIVELRYGKIFAVFRKMFFDEFGGEEGHFEIHTSGLREFKMNDILIRKACCWYTATYGKYGVHKGGIRFLSFPWILWDVLTSIKKQKTLQASRICSVKSPLADELSKIALFKCQNEEKAFVEFCNDIQKAVPIVKTYRENYGDHFLKACYVLNYWLLKERFYERTGMTSQQLVIIFLQFGIAIRHGTLNNYTGRIPRIFPKMITQSKDIIDIDSEGPLLPEIGLQIIEFLRYVSGYDFMFSECIDLSLGNILESTRLITRNTIWKSFSLASFKVFHHVTLSYSFRALHCGSTDVGNPEARQVAWDYGEIDNPLIVHKEALLVNTPGMPPVHLESTLNILQAWSGVDDLMVRPMNHRDLFIVTCAGGAESRQMLRRLLQLPPTVLREALLTDSIPSEIVTGTEM
>RSD-3 - L596_g5786.t2
MSCVIDSGMDFTFIPSTFAGRRSVGAGAMSDLFSGIANLTKSVTDTFNTYEIRKLGDKVQGYVMNYTEAENKVRDATNEDPWGPTGPQMQEIAHMTFQYDAFPEIMNMLWKRMLQENRAAWRRVYKSLILLNYLLKNGSERVVGNARDHVYEMRSLESYKNHDERGKDQGVNIRHRVKLIIELIQDEDLLREERKKAKSEGKEKYQGYSKEEMRMGKGGSYSSSSMGNIDDWNGRSYKSDNFSGGYRDEPSREVNSFNFPDDERNRSESPELGIRETKPVEDEDEFGEFTEARSTSNSIPSKSDGVIPPAIQGPGVAVHSPIKPPQGGVHLSLDNDPFGDFTSAPIPKTQPEVDLFGDFSTPAIDAIPNLPRPPTPGYSAPSNSNQPSSTLVDLFGDAVPSNPTPNAAAPQVDFFADFSNPAPQAPTSSVAANDFFASIPQMTNVTPNQSFVANFDSSVQTQAVSSNLDLFADVSLLSSPSHPSTNAVPPMASPMLFQSQSMTPSPAASSSHTPRSVASANPSKASSSMWDDMKGRVNIDLDNLSLRNSGNMKQSLSMNQMQKNKNQGPLF
>TSN-1 - L596_g28061.t1
MTDSAPQQTPAPAGAVKRGTVKQVLSGDALILQGPAVNGPPKEITVYLSNINVPRLAKRPAEGQPVSSDEPFAWEAREFLRQKVVGKTVSFVRDFTATSGREHGRIYLGGTSIENAENVNETGVAEGFFEVRTGKQIDEYAQKLLDLQEQAKSAKKGRWAFDEQQLKEKVRNVKWNIDDLRNLVDTYKHKPVKAVIEQIRDGSTVRAFLLPEFHYVTVMLSGVKAPAVRLGSEGRAEEYSEEAKFFVESRLLQREVEIILEGVSNNNFVGSVIHPKGNIAVLLLENGLAKCVDWSIGLATGGAPALRAAEKLAKDKKLRLWKHFKSVSSGDKKSFVAKVTEIGMGDSFFVQKDNGDEIKIFLASIRPPRNEAGQEKQSVGRQFRPLYDIPYMFEAREFLRKRLIGKKVNVCVDYVQPKSEQFPEKTCCTVTVGGQNVAEGLVSHGLAKVVRHRGDDENRSSHYDALLAAEAKAETGKKGMFAEGDPNEKGGVIRVQELTGDANRSKQFMPYLQRSTRPEGVVEFVSSGSRCRVYVPKETCIITFLLGGITCPRTARPGPGGKLIGESEPYAEEAMKFTRSKCLQHEVQIEVETMDKAGGFVGYMFVPNDKGGHNNLSELLVENGLASVHFTAERSHYYNQLNAAEERARRARLGIWKDFKEEKQIDALEQENAENVERKLNYKQVAVTEVAKDLLRFACQSYEEGPKIVQLMRDLQQEMNANAIAGSYTPRRNELAAAKFSQDKQWHRVRVEGAKAGMVDVYYIDFGNRETLPVDQMAALPSKFVTQAPGAQEYQLALVGVPNDPHYAAETIAAFENLVFSNSNILLNVEYKAGNTEYATLSIDADGTKSDIGKTLVMEGNALAEQRREKRLQTLVSEYTEAEQKARRARKNIWEYGDFTGSEV
>UNC-130 - L596_g11813.t1
MRFSMDSILCPPSATPDNKLVAASRKRLLTQSLTPSPPPVESKMAKLDSEDSSSSTSSEDRQPPAVTQIVVAAIDKGELAAAAVLRSDGDSSGGEELATEAKQNALAAAAVVGQHARGGGEDTTTSPGTSRSPVTSDEGDSGDECNESSNGDKRIGMSSTRSRSGATKPAYSYIALIAMAIYNSPEKKLTLSQICDYICNRFQYYRDKFPAWQNSIRHNLSLNDCFTKIPREPGNPGKGNYWSLDPNAEDMFDNGSFLRRRKRFKRQSSNNDFASLPFAPPGAHFLPPQAAFLANPAFVLRSPMMPRVGPHLGLPPQLYRTPYSPGFLLPPVTSTNGLPHGLPLSLSSLPPGMDHQRLLAAMAAQSAALSSASSPPNTVSPPSPAGAQAH
>XRN-1 - L596_g18210.t1
MGVPKFFRFISERYAALMERVQENQIPEFDNLYLDMNGIIHNCSHPNDDDVSFRISEEEIFGNIFQYLDQLFSIIGPKKVFFMAVDGVAPRAKMNQQRARRFMSARNAEHQQKQALAAGKPLPTSDRFDSNCITPGTSFMIELQKQLEFFIQMKQSTDAKWRDVRVYLSGHNVPGEGEHKIMDFIRTERSKPDYDPNTRHCCYGLDADLIILGLCSHEPHFALLREEVTFNRAGQKKDKVGIEGTKFFLLHLSLMREYLAMEFQDLKETTTYRKSNTDSLPFALNEENVIDDWVLMTFLIGNDFLPHLPNVHIHEDALPRLYKAYKAVLPSLGGYINERGILNLKRLETFFERFSVIDRDNYLDQFEDADWMRNKKQREQGEGPPDVIPQVVFEGLLEEEDVIESSSTCVSSSDIGAFDTDSEDERKEVEKITGVKKKKNGGDVIPSASEALAAGGWDSESSLEIDGLNLEDSDESPDENWNIVIHRSFKKKRRDYYAEKLNYVNISAEELDDQARGYVRALQWNLHYYYHGCMSWSWYYPHHYAPYLTDVRNFGDMKIEFDLGEPFNPYEQLLAVLPAASCRCVPPALRPLMTESSSPISAFYPTDFKTDLNGKRNDWEAVVLVPFIDENLLLRTAKAAYPSLTEAERRANTHTGDLLYTYSTKDLGELNSSCAKFDRVAENHSKCENMEKNAFRLPRSKIVTGLLPQTKLDVYFPGFPTTKHLNYTGSLEIAPVKVFHMPTRKPVMILKIGDNTENKYSASGVDPRSLLGQEVQVNWPLLKLAKVCSVVTPDGRYLMENHQTSFTDFGPNKHKVFKDIKDMVTDREFGRYGICVSNVKAIVEVNLFTGSRMKFKGNNVIVEKSWSDDVFAVSDGLVLQNVEVKNDMEQKFKSAADAFPVGSKVFLNSFKVSAYGMMGTVARNDVAARGTCLVEGVAPLQVNVKAAVSKKQYKKFWFSVFDLAKIIGQDKHVVNRVTGTVIVQMDSDFKDEEQEERGKHHHKHGHERPRNNINVGINLKYTKRNEAIVDFSKRENETWYYSLFTLKVVQDYAQRFPEVFRALRQNKDQYLLEDFWPELSPEERTNKAKEVVEFLKALPCSSRRPETCNYKYADPEELLLIQKEIAATVAETNVQQFTVHARALFRPDFVTGDSPPVPNVEFKILDRVIVAKTHRYVSPGAAGTVIGIKSQIGKETELDVMFDVPIVGGSNERTGPNGLKYTRVFSHQLLNITHAARLRGGDQSESPSDNQPQSSRQRFGNNRTDQRPQQHHKPGHYKGADSRMDNPKPSHHRAGNSRPENQRSGPPKIQVLQRPKQGDRQHASDPPGSVGQRLFQNSQNNRRGQNNRHQQNRQCQPKPERNEKAKSVEDQANKGVED
>XRN-2 - L596_g10155.t1
MGVPAFFRWLSRKYPSIIVNATEERPTEVNGVNVPVNSTKPNPNYQEFDNLYLDMNGIIHPCTHPEDRPAPKTEDEMFALIFEYIDRMFAIVRPRRLLYMAIDGVAPRAKMNQQRSRRFRSSKEAAEKEEQIRLVRERLEAEGVPVPPEKSPEDKFDSNCITPGTPFMARLADALRYYINMRITNDAAWAKIEVILSDANVPGEGEHKIMDYIRHQRAQPNHDPNTVHCLCGADADLIMLGLATHEANFNIIREEFVPNQPRPCELCGQYGHELDECQGLAREEPGPDQCEPMSKSTNFIFIRLPVLREYLEKELEMPNLPFPFDLERVIDDWVFMCFFVGNDFLPHLPSLEIREGAIDRLIKLYKDTCFTTGGFLTENGTVAIERAQQILNGLGDVEDQIFQERQRKEVQFKERNKAKRRRERQQAPAYFPKVGLLAPMSTPQTFSGERTREMARDARAEAMDFTNQQQRMQSIMQPVGTGNQPQIGQKRKAVDPAESSEEEEVHDEVRLYEQGWRERYYASKFDVSPSDMDFRRKVAEAYMEGLCWVLRYYYQGCASWDWFYPYHYAPFASDFDRIYDFKPDFSKPTAPFKPLEQLMCVFPAASRKHIPETWHHLMTEIDSPIIDFYPNDFQIDLNGKKFAWQGVALLPFVDEDRLLGTLSEVQDKLSEEEKHRNSLGPNRIFIGTQHPAYQLFEEIYNHEGGADVHIDSALAYGMGGSIAKDSLVVMPGTPYGSAVSHETCHDIESNACISAIYADPAYPEGFVFPAVRLPGCKEVEKTLKPGDWNQHRNGNYRPQIGFDRHAPRASLDQGAHRGFRNEVRDRSFNNDRRQSDRFDRRNDRHPYRGQGSPQQHHNRGPRHSYPPPGGHGGHGGGYQHGYPQRGHPHDHWNGGRGGRGGRGGRH
>VIG-1 - L596_g22586.t1
MEYGINVNNKYGYLSDEEAEDPEVFIKKAIQRKDQKMAEEKKMAEEKAAAEKAAEAATEQKRAANKENRRETTERGTRGARGPRGPRGEGFRGLRRDGPKPEGEGTGPRGPRAPRGEGRGGQNRPRRERPVKEGETEAAVPAGGDEARLNEEGDNRRGGARGGRVFRRGGPRVPGTRLDRKSGSDQTGVRSIEKKDGHGKGNWGTEQDELAAAEDVNVSSGGEVEREKTEEDLKREAELEKAARELTLSEFKARMAQKADKPEFNIRQAGEGVDAKNQPKLVPLQRENDEDHVAEEVVVV RREPKNKRLNIDINFGDENRQRGGGRGGARGGARGAGRPTKERRGKDTHFDFTAESFPALGAR
~~~

## Argonaute proteins

~~~
>ALG-1 #1 - L596_g7718.t1
MTTMTGGPQQQQLDQTQQQVISMLDQLSFSEGGGMPPNYGMLGGPLMGPGPLAPGPQQMPFGPGPPGPQMPQDIYFQQFAPPGMPGQMMMGGGGGFDPRPGSLAPGAPIDPSQTMMPSAMTPSLVSTQGQGAPGTPSQLAPAIPASTIFQCPRRPNHGVEGRAIVLRANHFSVRMPGGTIQHYHVEVQPDKCPRRVNREIISTMIRSYTRIFDNIRPVYDGKRNMYTRHPLPVGRDRVDLEVTLPGDSAVERKFVVGIRWVTTVNLTTLEEAMEGRVRQVPFESVQAMDVILRHLPSLKYTPVGRSFFSSPTAGGAAPAQGHFQQESKLGGGREVWFGFHQSVRPSQWKMMLNIDVSATAFYRSMPVIEFIAEVLELPVQALAEKRALSDAQRVKFTKEIRGLKIEITHCGQMRRKYRVCNVTRRPAQTQTFPLMLESGQTIECTVAKYFYDKYKIQLKYPHLPCLQVGQEQKHTYLPPEVCNIVPGQRCIKKLTDTQTSTMIKATARSAPEREREISNLVRHAEFNADPYAHEFGIAINTAMTEVKGRVLNAPKLVYGGRTRQTALPNQGVWDMRGKQFHTGIEVRIWAIACFAQQQHVKENDLRTFTSQLQRISADAGMPIIGQPCFCKYAVGVDQVEPMFKYLKQTFAGIQLVCVVLPGKTPVYAEVKRVGDTVLGLATQCVQAKNVIKTTPQTLSNLCLKMNVKLGGVNSILMPQVRPRIFNEPVIFLGCDITHPPAGDSRKPSIAAVVGSMDAHPSRYAATVRVQQHRQEIIQELTYMVRELLVQFYRNTRFKPTRIIVYRDGVSEGQFYNVLQNELRSMREACMMLERGYQPGITFIAVQKRHHTRLFAVDKKDQVGKANNIPPGTTVDVGITHPTEFDFYLCSHAGIQGTSRPSHYHVLWDDNNLSADELQQLTYQMCHTYVRCTRSVSIPAPAYYAHLVAFRARYHLVDREHDSGEGSQPSGTSEDTTLSNMARAVQVHPDANQVMYFA
>ALG-1 #2 - L596_g21728.t1
MQQPDEKKPMKPIPLRAMEGSESVQVGRLNVQVNGYTLRIAADTQRTVVHQHEITLFGVFSRDEGQKDVNLLNMGGGLKDYKKQGRRLIIYAVFDKIVEANPLIFPKNIYQCVYDGGNILFCKQQLVDYKLPMESVMESSKFCGSVRDFLGARCEKIKFKITYTGVVPLDTKELNGNSRSLVQFLDIVTSQQICRSEEQLVFKNRRYDSDSVRDVQASMAKIIKAGSEKTISIVGENSSHQEALLLIEPKRSPFFMGGNLKDIFDIVQREMGVGNPRLVSELTKLVKGLGVFTLHAKKLRQFQIKDLTKEPAGRQIITIERSGQATDLTVAQYFQDMYQMAVNPNLPCIACEAIIRGQKQVLLYPSEVLSVMPGQRVQTQKQTPKLVEELIRQAQFVPADLMQEVQKERLLFGLENSKYLAEFGIKLDGKPREAPAKVLPTPAICYGRDTTVQPDQEGRWMIRENQYFKPATQCKWALCVVENAMDSQVATRFKDSLVRAAAGHGLMIGEPSLHRFQKADPEDLTREFAYLKANKVQFVMGIFGGDRNCIERNLLKEMEIRFQLITQTIQSKTAFKGTTNKMVIDNILMKTNLKLGGLNHQVTTSRAYATRFLDAIFPKNRIFIGLDMQSPGSPMLGGINEFTTDPTIVGMCVSVKNAAQMRGHYWCQPATFKFIMGIEKALGSVLKMYQAQRENVNDDDFPTDIVVYRGGVSDGDIPMIVDEEIPQMKAAFANLKIRGRSYCPHLTVLLAQRASERLMPVSSPMDGGGGYNNPKSNGNVAPGTCVSSGIVSPTRSEFILAAHKAIKGTAKPMRYIVLDEYGSQSKRFTIQELENMTNQLCYTHGIVTSPVSRPGPLYGATDLVKRGRADWKAREYRSRNAAPVGPVDDQFLEGYNKQRLAWQLEYDGKFWA
>ALG-2 - L596_g16709.t1
MNVFLSTAATQLTIPIDRATDRTGGVSHSPPAILRRIFWDVVDNNRSLFPAFGIVFNDKDQLWSKRKFNQFEFEAQFGSNNDHTIKLKLVDMFDFQISPTSDQIQAEFLNSLLTQTDRCKLLSVLCLDCPTAISVCTCAVFLKMRIPIIDYFLSVVSRRPRLTDADYRGLENFHWNPDQITALRESLKGLTMVTTYGGTSYYHYKFLDVYRGNASETMFNWRRPENNEEEEITIQEYYFKKWGIRLRLPHFPLIQAAPLQKNIYLPMELLMISDRPQRFQKPIPEDCMRAALEKATISPRDRFDLICDMIQQSSINDDNRFMQKFGVYVDTTLMITEARVLLLPILTVRDGHTVIRPDTTATWRSREVQSNSRRRAIIAVIINSEVRSSMGNSFFPYFNALMRACRSIGINLVDQQDSFQPVIHTYNRDQQEIDVAVXXXXILSFSGFTIPNSVKFFVQRDRLFFIPTEGSRPGADDLGRGLQMWYALHSLVSVGEKAEAIVNYDRTCAVFLKMRIPIIDYFLSVVSRRPRLTDADYRGLENFHWSPDQIAALRESLKGLTMVTTYGGTSYYLYKFLDVYRDNASETMFNWRRPEDNEEEEITIQEYYFKKWHPTPTALSSDSSRSPAEEHLFADGASDDIRPSAVPGIKQ
>ALG-3 (T22B3.2) - L596_g3457.t1
MIDNIDERVSTPIQVIDIIFPSTSSLTCPLISGATSTESDSASSSSRLSTTDDSGSFVSDSEEHLTRPSLSPAPSCPQLPKLGAGSRAPSEGSGRFMDGESSESEDEDEAEAQQRRVIAQKEAGCGGLLSEDQSKMQIRALMARPGFGTNGRKIPVLANFFEIGIRNKDMIVMQYHVDITHPGNRKLDRDENRTVFWKAVEQNPQVFQNRFAIAFDGAHQLYAVQKLRLPHGGSSAEIPIDIALARDLRSSSKCAINLQLVGPMIVDIGKSKSMNIDERVSTPIQVIDIIFRQSLTCPLIANSSNFCAWKSSFYRLPTPNSNDALDLEGGKQMWTGFFSSAHVAQNYRPLLNIDVSHSAFYKQHIEMVDFMCEVINERASAFTCRATNPGPGAYRGGMKPGGPRGMGGNINDNSPGFLSRDKLYENFSLSSQELKVLDDAIRGVKIRVTHRPGVVRVYRVNGLQVSADQLTFVGKDSDGAENRLTVAKYFELKYAKLKYPRLPCLHVGPPSRNIFFPMEVCRLDSPQKYAKKLSERQTSSIIRAAAVDAEQREKRIVSLVQQAGFDTDPFLKEFGLKISPQMVETVGRVLRPPAIQYGENNRRMDPIVMPKDGAWSMDNQVLYLPAACRSYSLIALVNPREQPNIQNFCQALHQKAQNMGLMLPQWPDLVKYGRTKECIVQLFKEIAFEYEQTKQQCDLIIVVLQAKNSDLYMTVKECGDMNYGIMSQCVLMKNVQRPSPATCSNIILKINAKLGGINSRVVPDAVTRKFLIDVPTLIVGVDVTHPTQSEERQNIPSVAAIVGNVDCYPQTYGAHVKVQRKCRESVVYLVDAVKERLLCFYRNTRKKPARIIVYRDGVSEGQFGEVLREELIGIRKACMELSSEYRPPITYIIVQKRHHARMFCKNPRDSVGKAKNIPPGTIIDTGVVSPEGFDFYLCSHFGIQGTSRPARYHVLWDDSKFSADELQQITFSICHTYARCARSVSIPAPVYYADLVATRARCHIKRKLGVHETEHFDMDKFSRKPSTTTTTTTPFDNEIVGARGDCRKGQMPDFTNLKQSTCEAALQDFVNVTEGFKTRMYFI
>ALG-3 (T22B3.2) - L596_g16708.t1
MTTIAGYGDQSVNLRAAIQTSLKEFYKGTGGLYPSHIIMYRLGCTSSQMTAAAKNELRAVFSAVKEMAELANQGHFDPTITYLHIERKHQKRFHCGADVGNLPNGFAKANNGNVPAGTIVDRDVTSQAFYDFYLCSHHGALGTSRPTRYIVFYDDWNLTADQIQAATFCLCFLNSNCTKAISLPAPAYYASKGVERGKKYLNTCMLRREDIQPNEPLMMPKSISNSMYII
>CSR-1 - L596_g20174.t1
MIGEVTLADLHAESFGAPLQYPDLPCVEWERQGPARFFALETLIVMPDQRVAFERTDARQSAQLQKINTVRPEHRLKNIEDQMRRLHLWGDSQCPVLTGFGISIEQQLLSITAGVRTAPVIQFGSSTDKLQPGRPKWEMALKKNRYRIPANVKSWALIYRCEDKDVDVVRLFTTKLRSTAKMRGMNVEVPLAVIRFSPNRQAGEGDDEVLGRHVNEARSAGAEFIVYVSSKCIQDHDLLKLLERRRLRVTQHISLESVRAVVIENKITTLDNIIHKLNVKNGGFNYTPLIEHVGNSRELELASGNVLVIGLDVAHPAPMNASQRRMMHSVDATIRSLEPSSIGIVANVIKNPHAFVGDYHFQTARREAVEPRILKERVKWIFDLLAQNRPEHQRPKHVVYLRDGVSEGQYTMTIRDELGAIREAVREIDPKYRPKFALIIVTKRHNKRFFDSSKGVVGNPLPGTVVDHSVVRTDITEFFLQSHIPILGTVKIPQYDIPVNEGGFFMDELQAFANCLCHSHQIICTPVSLPEPVYAAHEVAKRGHNNFLEFRRSHPEMVPYVDGNVNIIDCEEVTAKLAYRGTPLEAIRFNA
>C04F12.1 - L596_g11107.t1
MMGSKYPDKLPAHTKGNPKVPLVSNSFEVSLGDKIIHVYDVSIIQDCESRGRKKVIDWSTSSGDSAKRRIKLAVSKEIFKKALEVKKFASKEAALVFDFSKILFSSEKLKDHLCSAIILTPEDFDVLPSFEGNSKLCRGTYTVRIVPTKAGSHQFRANDLEKALAKNQEDHSLRQFLEVATSFKPIQEGTHKFFRGILHDVRTSAPGRRDLQNGPFEIAYGISKGARIIGDMDNPKAALVMDSKRFAVYSESKETFLEDIQKVLGDKDLKIHLPRITKMFQGVTLGHCFNKAVEVKFSSLSNVLAEKLTFENSDGQKSFIVDYLEERYENYQCRARRWPVVVDKFPGRSGDVCYYPLDILYVKEGQLVPLPLQQEFGITQELLKEVSKPHLRSAEIGRAPKDLELNARNAHLREHGINVKQAPIKLEGYRAQPPKLGYANGQTAAVDANRANWEAGRYIYPAKVDSFRYFVRQGCMGRDQANLFLNKFLDMCRSKGLDMPKPQIEFIKGPLALKNLLSAEDKAVGKKKSVTFVLFVDSEKSKTHDALKFYEAKYQILTQQVRSETTFKAGRQTMENIVAKTNEKCFGQNYAIHGDEFISTKDTLILAYDVWHPTGASAQKRILDIPDDTPSVVGMSFNGGVHADGFIGFYAYQEPLQERVDVLKSYMQHILRIFKKTRGLLPKNIVVIRDGVSEGQFDMVCQHELASIRAGCRQFANAEKVGWNPKFMVVTVTKRHDKRFFVQDGHRVLNPPPGTVVDCTVTRPDMTEIFLQPHRPFQGSAKAAAYSLLVNELEIPKQKSGSDLWLTNFLMKLCYSHQIAPSSISIPEPVKQADEWAKRGAANLEFLKRENGDKSLDMHNFLKSSSEGSFYDWAALSDALGYHTKRLEGTRANA
>HRDE-1 (WAGO-9) - L596_g11197.t1
MTSTSVNLNSFKIDISETVHRVQQYELVFVLYKSFKPGFTATPSRTRVPLGFKYKYNLDTFQNGDHWVGGYDIAYGPQNPVKIRMRKEMLFHFFEFFKQQFSHTLAIGDVVPMMIFDCDRTIYSNQQLKVGRGFSHTWSYIEHLPEKVQDHIDILCGNVSDFCGLSVFLNRVGEVDMTDLGSEKNPNKGVIQFLETLLWQPLYTQFTDHVIYGKTFFDLAEKNRISVMQDGLFLATGYDISVDLIPDYEKDMSKLMPVVRFTPGSSVFYQNYSRMDALLMSLSGTNGRDTLYQLEYDCKHPSIVEKLSQTVRGATVQFIYDSDQIFEVDHLDPRTPHQITFLLQGRRRISVTDYFMAKYNIQITSELPCVARKTRYGMSFYPVEVLRVAPNTKAIHKYLPADVKDMIEKGTILLPSAVQQRIKEAIGEMKLYSAGIRKHDMARINPFLFVFGIALVDKDRPVRIAAHVSESPLIEYCYRRGATSTLMKTEQNHPGVWNKSMKFLPPFLKPADLGKPVKIRFVNTFVEEVHLINRFMDKTIHKLRLKGIDVMKERTISGSERFGKIRDARDIFGRNKDYRTMLKAAEDILTSSGADLFYIIGSTDPMNPTRDIFKLAEITKPSDKRIVTQHIGCKTLCTTVSTGLSKRSPGILESIVMKTNMKLGGTNYTLREKDRSSGYRYGYPRSTHLIIAFDKVNPQHTDEKGEKLYHPKACAMTYMIPHTTGLITRGTYWFQTSNETNLTMMISAFEEALKFYHKDSRGYDPEKVVVYWDVSHGEKEVTQEMDAMMAIVAEQRGESVRGPFLTFITVDQNHNTLLLPTEANVRDRLSLQNVTAGTWIQESTIGESFTMVSYESDDKMTRAVKYDVKVAETHMHSIQDLTHKLTYLQNSGWRSVSVPAPLKGATKLAKRAMKSYDIMDQIRGRSGPIEKQLFEERVKEISDWIAVKHLTNYWA
>PRG-1 - L596_g25491.t1
MDARTGQLKSFRAASRQNRGPQTDESLICFAYQNHPRFIVKRTKSISLLKTLILRRFDALASALMRQQKSIIYDFQMVIALKTKF
>T23B3.2 - T23B3.2
MRSRRIYSFNSSIEMSDGEIPIRVVDNRNRDDLLKILLLVVLIIVFPPAAVAVQANECNVHVWISLFLMLFFIIPSYIHAVWYVFIRKPKELTIA
>WAGO-1 (R06C7.1) #1 - L596_g19943.t1
MSDFENPNVKRPVTRIVDDVEELFERLQLEPGKPVMADKQLPGTTGRKVKLQTNIYGLSLQKIPVHRYDVNVIARLGEREVLFTKRSKEDAVSTDRKDKCRTAFELAVNTFGEQFFGQERYALYYDCQSILYSMKPIPALEDKMSHELSLLPHHLTDFPAFTGLDAVVVEIKKVADTFNLNLGNLGFLNRELIEQDHSLQQFLELITSQHALFTPADHICYGSGTSFLLNHKKYGFLDEDCPDLGEGKYLAVGSHKSVRFVEGPGGRSGAHAAVVVDTKKAAFHACENLIQKAMAIINLTPQTHCRKNEVDKLRGQLKGLFVHTKHGRRQRLFPIASITEDTAADKQFEDPDGNDVTLQIYFQQKYGITLRFPHAPLVVVQENKQTNYYPMEVCYVNDNQRVALNQQTPLQIQKMIRACATPPAQRVRQNKENMRALELNNSNRYLQAAEVRISNNALILEGRVLQPPDVYYGNGVKAVVHPDKGSWRLQNKPHFLLAVEIHRWAIYVVGTGNRDILDQPKFENFIRMYMAECRSRGIRINDPCEHRILPADPEIIKDRIKAACEGECYFMYFITSDAVTDIHKIMKYAERECGVITQDMRMSSANDVVAKGKRQTLENVVNKTNIKLGGINYDIRFNSPDLNVLRNDRLFLGFAMSHPAPQTQHERNKGVAPRSPSVIGFSANMKSSPVDFVGDCVFQEPRRDEKIGVIRGVVNNVVTRFRDSRGYLPKELIIYRNGSSEGQYPLILKYEVPLIKKALEEVQCDAKVTLIVSQKMHNVRLMLSQINERDRAPEQNIKPGTVLDVGAVHPVYNEFYLNSHVSLQGSAKTPRYTVLFDENNYNMDQLEYMTYHLSYGHQIVTLATSLPAPAYIASRYADRGRNLFNASNTNWNVLQDGQLDYNQITRDLSYGVSDLRDYRVNA
>WAGO-1 (R06C7.1) #2 - L596_g995.t1
MDALKGHCESDETLYKIETALVSKSGTIRFYPQEAKKVLNWAVEPVMVEEDPYKEEDGAFYSLCHVNIEFLASGVHCIAKLLAPFSNFRRSSATKSSSRRPKTTANVENRSATEPMTNLRSKIAGPATKTASITVYLTMESVTTKMAHLTMAPKLGAGTGGRPVPLVTNMYQAKMMRAQPVYRYDVAMEMRFGAKSVSLVKKTIDDMVAIDHKDKCRAAFRIAVRLHDKVLGSPVGLFYDLQSTLYAIDKIKDAKENTECKEIELCIPSDMLKNNDYFKDRNPDSVTITIKRVEADYQLSLGDLSFATDASKAHLNADLVQFLEIATTQYAYLKAGDVLTYSSGLIYFSEKRRKLNGGNLLIDGVQKSIKVIEGSQKGQPQLAVVLDPKKTAFHAKDVAVSDKIYEAGFMEDDGRVHPAKLETVKNQVKNLFVEVRYGKKPTRFMINGIDKESARVKTFLSSTGETTVESYYLKQYQIELQYPLAPLLAANKKVNGERTTIYFPMELCYVCDTQRVKNTQQTSKQISDMIRSCAMLPADRVKEIKDCAKRLQLNGDAVSGSLRSAGLSVATQLTSVQGRALPPPEIVYKDNKKFSVDPNSGKWKATGAQKPKFLLGASINRWAMMCVADRPQPRDDALMKNFAEKMVRECQGRGMRVNNPVFYQAVQGRPDSLESMFRRAKQDNMEFLFFVQDGRLQAHKDIKFFERKYEIITQDLNQQTCRSVVEQNKFLSLENIVNKTNVKLGGLNYSLIVNAPNTQHLFAKGRLYLGFQVSHPAPLSDDQKAKGMKPKQPTVVGVAGNITNQPAAFVGDVFFQEPRDDRMADAMETLVRDFALRYKDAVGVAPAEVIIYRNGASDGQYQTILDVEVQEVIRAALTAAGAGSAKLTYMVVSKLHNVRLMPAQITGIKAPEQNVKPGTVVDTNIVDPVFAEFYLNSHQTLQGSAKTPKYTVLYDQNNFPMSYLELMTYVLSYGHQIVGLPTSLPTPVYVAGRYAERGATLLSASRHDSDMCKFSELTEALGYGSTKIGKNRLNA
>WAGO-1 (R06C7.1) #3 - L596_g15757.t1
MDTRSQAQGGFDTSIKVVYEMAPKQYAPENKAPVDLVCNAYRLRMPQPSSDYPARVYVYDVTLTLTRADGSLLTLVKTKMDDYTHYVSKQRCRSALEAFSQKFPGFFRTNEQRLFYDLQSILFTRVELPMRNLTEAMSIEASEFSRFGFAEGGQGMQRLNIVVQKTQSGTSIGLTDFSFLSSDIAEVGQRHDLSQFIEICSSQNAYMNPDQRVTFPGGVSYERGNPGDLVDGSKRVWNGVRKSVRYLTGAGNPVPAVVLDARKSAFHKEGEMVSDKVYAMGLMNDDGSVFDYNIDNITRQLKGLFVMVKHLNNQRTFPILKLDKKTPATYRFTSDDGVEMSVADYMSKKYGTALEYPKSPLVVVWMKQREVHYPMEMLYVCPNQRVTTNQQTSKLVSEMIKKCAIVPSERIKEIKHQATSLHLHDGALREVCIEVDPNMLHLQGKAVEAPKLKYGSSALVSVERNTGKWRTSNTKFLHPVQIKKWAVIVLPTGGKLTGQDSQVPTKFVDLMIKALIIKGVQVGPPAYTGMSSGCQLEEDFKECARDGYDFMFFIQDSKLQFHKDIKVMERRYEIVTQDLNLNTARNVVQQRKHLSLENIICKTNVKLGGVNYSIHIDRPEFADFFHDRRLYVGMQMSNSRVFELGDGSDVEKAGKPTIIGISANVDRELSSFVGDFMCGPANIEDLSSVVTEIFKYYSEEFLKMRGHFPEEIVVYIGGISDGDMPKLLRWHIPAIRHGLNTAKCTAKMTVIFTSKSHNVRLFPKNVTGERAPEQNVKPGTVVDTGIVHPEFTEFFVTSHQTLQGTAKVPKYTVMIDDNNLSLSYLETMTYVLAFGHQIVALPTSLPTPLYVAGRYGDRGGVLYNAFDGKDDLYAVNDQLTFATSKKLCGKRVNA
>WAGO-1 (R06C7.1) #4 - L596_g16574.t1
MPTGEDPVLAEPVASTEAEDNASPRTPHAIHEAGDAIGLERRLEASTSARDGNFYVALGLPILQGVVGNKKEALAVGVPIELSTNAFGFVLPALPVWQYEIEIEGLLAGSERRVFFTKRSPDDAFKIRKSQECRRLFQAIKHKYAHSFSEDNETYFYDNQGLLFAIAPLDLSLNEKMDCMLTAEELAALSPPTYASFSEVIVTIKNVAANPMTVGHVMTYTSQVLENNSRRLQQFLEVLTSQHMLANPNEFLTYGTRSAYLLDTSAHGLEAGHLTKDKEIKIGCEKSVKLVEGPMRGGQPDGKAVALIDVKKTPFHIPEGTVLDKARAILNREPKPSDAPRLKKDLAELVVYTKHTSKEHRYVVENVIADTAVSMTFPWTEEGREVTLSEFFVQKYRQNISFPRTPLLVARFGRERKLIHLPMELCYVARNQRVTSRQQEVDNISAKMIKACAIAPAERQLQIQETVNALQITSSNPYLRAARTKITAAPLIVTGHRLEAPKIAYANNEVLSPDARFGFWKPPNAHRRPKFFKPAVINSWAIVVLPSQAEFLQGDIISREILARFTDLFRSECRDRGMQIGQPVFTEFMKADVQQLRDLIKSLTRPDPSVCRPPLRYVIFITNGGITFCHQPMKYFERETEIITQDLKMQTVVNVVQQNKRLTLENIVNKANIKNGGINYVVIRNMPGQRPILKPGRLVIGLAMSYSVRRQAEEISTLPTAVGWAANITREEGELIGDFLLQESFKKDRVAVIQTIVDRVADAFKHPGGPKEVILYRSGEEGRFRAILEEDLAVLRATFDNMASKPKLTVIAVQKHHNLRLMPTKINRQDRPSLQNLVPGTVVDRYVTHPTFTEFYLNSHVAIQGTARTPKYTVLQDDANMSLEELEEMTFGLCFNHQIVSLPTSLPSPLYIAGRYAERGMTLYRQHQEHEEDQGTSASSPSSGSSGEPHLDVERLGTHISYGSSRKLKHLRVNA
>WAGO-1 (R06C7.1) #5 - L596_g20961.t1
MEREGTELRPSTIVRQTQNAAASSAPAAQVSTSAGAPPAAFGIRGPATSAETSFYSQLVNIPTPPLEEKKQPGTTGREQFKLRTNVFGLSLPKDAQVFRYSVDASGTLQRNDRRIEFAKRVGDDITYLNRREKCRHVVDQVVAKNPAIFGDRRELFWYDSQSILFSRNQLDIGSEAQFVLDQSDIGQNTLFEGFAHLKMVIRPAQTNFAVSIGDLEAYIQAELFESDHALQQFLEILTAQYAFNTPTEAMSFGTRTAYLLQPEKYGFKPADCADVGDGKFLGVGCDKSVRFIEGPGGAGGQRAALVVDLKKTAFHKVQSLYEKAREILNNRDPKSTDASRLRMQLKGIVVETKHGSRRQEFAVDNVVADTPATKKFKDLTGHEVTLQQYFQQKYNITLQHPDSPIVLTDRTKKFAAFPMEVCWVVDGQRVALAQQTPVQIQKMIRQCAVPPADRQRQILGLVQGLQLNSENKYHKAAGVGITPTALQVQARLLQNPIIVYGGKSTMRPDEKATWRLARQKPVYLKPVKIDKWAMFVIRGGNRSDCVDQAILNQFSNMMVQECRARGMTVPDRPTGLSFIGASREEVQETLEKAKTEGNQFCFFITNNDVTHIHQFMKFQERKLSIVTQDLKMSSAFDVVKKGKRQTLENVVNKTNIKNGGINYSLRFDDPVFNMDKLLPKDRMVIGLSTTHPKPIPGKKEQDQAPQDKKKQMHEQRTGPPVPSVVGVAANVLTESIEIVGDCLFQQPNREEKIALLQPVIRSLMLQFMKHRGMPPVEIVVYRQGTSEGQFRNVMELEYKMVKAAALQQGLNPKITFIVVQKMHNVRLMPTDCKAGDRAPEQNVKPGTVVDTMVTHPKYNEFFLNSHVALQGSARTPRFTVLYDENRLPMDEIEALSHSLAFGHQIVNLTTSLPAPLYIANRYAERGHNIFIASQEDYTKSKSSLQSPHSTTIEDNLDFSRMMNELSYCNSELRDKRVNA
>WAGO-1 (R06C7.1) #6 - L596_g322.t1
MSEELHAASKLKPTEGIASRKVKLTTNHFTLSFKSSKPVYRYDVAMVHYMMTKDGEKSRDMCKGERDDAAILERQRRCLVLMEAAKKVAHFTPSKSCCVYDNSKTLFSSEKLLENQCAQIRIEGEHIPDGFKNHPKLKQGYFFIEITPVSTNHKFTIDDLKSGVSDDLLNTDHTLRQFYEILTNAYAVTNDSHMVFYGNLYDNDKGESGRKKLKEARNLIFGVNKGARIIEGSSSRLVAALVLDSKKSTFFDDANNKGLAGNIRDLLGSHFNAQPHEVVIGDHNRKTVVTYLKDLRVYCRYQEDRDFVIAGVTKEPIEDLYFEYGSTKTSVMDYHKQAYNTKILYPHWPAVIMQGPRSKNYFPIEVLGVSKGQRVPISKQTPGQMAETIHDCAARPHIRFAEILKKLDGLNLASNVPNEFLQSFGVKIDCNPIQVQAHRRLAPKMVYAGNKEVDYDDIKGNWFANNAYILPAKIPKWFVITDRIDVSIVRKFAGILKDTMKMKRMTVGEPQFLEMPVAQLDGFLGKMAKELKPEDQSPFILFADTNDDSHALLKLYEAKHQILTQHLRARTVIECLEPRKRLTVGNICNKLNCKNYGLVYAVNPQDHAKTMYLSKGDVMVVGYDVSHPEPQPAHERRLGIPPTTPSVVGFSFNGGVHPGMFIGDYQFCAPRQERVDILEERIQWMLRVFTTNRKTLPSRIVIVRDGVSEGQMPMVLQHELESIRRGVKKLKAGYNPKFLLVTTTKRHAKRFFAETERGIDNPMPLSVIDHTVVRPDVTEFFMQAHKAIKGTAKMPAYTLLLNELGMTLDQIQSFMMGLCFEHQIVNSPISIPEPVYQSDEWAKRGHSNILAFFRLMDCPDPQKPSQKMIQRYLKASENPTTGQPEVEGYDWVRISKMLCYRGRRLEKTRANA
>WAGO-1 (R06C7.1) #7 - L596_g9675.t1
MLEIRIPPIVDDQKRRSETELNTVNVRTNVYEFELPTGTSEIYHYEVAVIGKLKSGRTVDLTARTTNDVLTLERRDSCRKVLGIVSERYPAVFDHNLVFYDQARLLFTSKNVEINTAAGIYSVILGCQDHLKGDEMFAAFTSVEFRLARVSVVEVGNVQKYLNKNLRLVNHSLEQYLNVLTSQHVANKTITFGNESVYLEQDDQCGVRNNVDLGSGKLLKTGFAKASRLIEGLTEQPAAALVVDLKKAAFHVQQTVLDKARLILQADRKRCGGSQPRIEDTEDLSRELVGLVVETKHGSRVRKYKIAGFDKSTPASRSFEKDGRNVLIADYFAQQYKVKVENLDTPLVIVRGYGRDFHLPMELCYVKDQRVGLKQQTPDQIKDMIKQCAVKPVVRIAEIEKIVKGLRLIDERKDILGSKVGIKDKPLKVEGHLLPPPLIVYADKDTCNQRVPPSREPASNGTWSSLTANTTPAAFFKPAKIEKWAVIAIHTEDNGDPHDERNKRILAQNTLETNILHQFATVFTEECRNRGMTLPEAEHVKFLDDTSTTVRDFIHQAKLSFVLFVCNNAITHIHQSMKCYERKFETVTQDVRMATVNDIMTKRRFQTLENIVAKTNVKNGGLNYNVEMPSKNGVRELMPKGRLVIGLTVRVVRVPKSPKEIEEDKKKAEEREKQHKKARDSKFARKESTTAKLKPIMTVGYAANFTDVPTEFIGDHLYQEYRETGDILGMQIIFERVVLEFKRARGMFPTEIVIYRECTENSDFIGQLQLEQMLLKSAIKSTASFRPGFEPKLTLIAVQKRHHVRLMPIAMRREAKAPEQNLQPGTVVDRMITHPEFTEFYLNSHTTLQGTARIPRYTVLKDENNLSMHEVQLMTYGLSYAHQIVNSPTSLPTPVYVAVTCAERGMNLAKHNFRAMNIKFNANDDYSKLEDQHVDISHLNEQLSFGNCKLSSIRNNA
>WAGO-1 (R06C7.1) #8 - L596_g18735.t1
METLTKAVANMMPPVNLASKREDAKPNAQERALPLQTNMFSLSMRDEVPVFMYSVDVFMKVRSKAISLVKHSRDDYIVIDRKNKCRAAFRFVVRANPAVFGNPGKIYYDLQAQLFTLEKLNMDNEDEGLELVIDGADARRSTDFAEIPLDGIVVQIKRAGPKFDLALGELQLKFVAQEPKSHELLQFLEVATSQYAFLTPSDFVTYPAGLSFIKQKDPKAGTELEGGKLLLDGAQKSVRLIEGEKKTDGGGKLAVIVDAKKTAFHKTYQKVIEKVNDFGFLQSDGTVHRMRIPDLAKLLKHIYVESRYRKRTQRFLINDVNPDNARNKMFTRNGVMISVEQYYREVYNITLRYPLAPLIVSKPMKSKDSEEKMVCLFPMEVLFVCPDQRVKINQQTPRQISDMIKRCAMQPDVRVKETKNYAEKLQLNGPASQACLNYAGVSIANEVVKVSGRKLVAPEIQYKDKKIAVDSNTGTWRSSGREKPKFLVGSSLKKWAVYLLGVRQPLPDENLGRKFVMRMLEEARARGMQWESPTAVKGVLGHVDNIEKLFKAAQKEGLEFLFFIGDQKVQVHTELKFFERKYEVITQDLDLKTCRNVVEQGKFLTMENVIAKSNMKLGGVNYSIQVNDPTIKQFFKPGRLILGFQVSHAAPLKPDEIAAGVKPRVPTVVGVAGNVSKGDPACFVGDFFYQTPREDRMLEAMDTLIADFAQRYKRATGKDVAELIIYRNGASEAQYTDLVRDEIPSIRDALQSAGFRGVKLTVMVVTKLHNVRLMPVAIKTGAKASEQNIASGTVVDKGITHPKRAEFYLNGHVTLQGSAKTPKYTVLTDDQGFTLQQLERMTYALSYGHQIVSLPTSLPSPAYIAGRYAERGALIYQGFNHTVQHDADLQTLNGTLGYGKSKLGDKRVNA
>WAGO-1 (R06C7.1) #9 - L596_g18947.t1
MSNPSHNGGRRPGANGNGNGSQPGGLEMRAATHVRQIRSGASSGSMSSEVRYYQELTQAGQQLHLEEKKPPGTLCREELIVHTNVYGVGLPDVQVYRYDVDACGFLERNDRRVEFAKRAVDDVADTTRRNKCRAVVQLVCQNYANIFGNHREYFWYDSQSILFSKNMLDGIPEREKKEFVLTQEMLAGNPMFRGFKNVRLNIQRVSNSFAINIGDLTQYISADLEENDHSLQQFLEILTSQYAFNTPSEAICFGSRAAYLMQPDKHGFKAQDCADVGDGKYLGVGCSKSVRFIEGPRGDQRAGLVVDLKKTAFHYEQSLLAKSRAILNREPRGNDAGRLRQQLKGLVVETRHAQKPVRFTIEQVSDETPRNKKFMLQPDNREVTLLQYFQEKYNITLESPDSPIIVGDRLTKFAVFPLEACFVVDGQRVSMDQQTPTQIQKMIRACAVPPADRQRQILSLVQGLQLTSSNVYHQAAGITITEKPLLVRGRILPNPKIIYANNVVVSPDAQKATWRLDRQKPHYLIPAKIEKWAMYSIRAGGRSDVLDQPTLLRFAQTMVTECRMRGMQLGNPGEVSFIGCAQEEITATMEQAKKDGCTFCFFITNNDVTHIHQHMKLLERQTGVITQDLKMSSAVDVIQKNKRQTLENIINKTNMKNGGLNYTIRCEGLSNEQLLPSQRLIIGISTTHPKVAQTALEDRDRADKNEDPLTKKKPHHEKHHRSEHITPSVVGFAANFKKDPIEFIGDCLYQYPQRDEKIGVMQPLLRAVINEFSNNRGVPPNEIIVYRNGINSISTMELEFLMVKAVARCQGITPKVTMIACQKMHNMRLMPAKINPRDRAPDQNLKPGAVVDSNVTHPKFNEFYLNSHVCLQGSARTPRYTVLHDEGQFSMDELQALTYNLAFGHQIVNLTTSLPTPVYVAAAYADRGHNIYGVSQKDYTKSRTSDLNSSTLEGNLDFNRIMMDLSYSNSELRSKRVNA
>WAGO-1 (R06C7.1) #10 - L596_g1747.t1
MASVDVELREKTKIRQEADRMFKEDYGIDVGSTEMPAKMMKPAAQNTMSLVTNLFPVKTHQQMPIYRWDVDITIAGKNGKTFSLTKKSDSDAVAVNRKLKTRSIFKRLVAAHPEKFGKLEENYYDLESMLFTLKDLSTDQEAQDFVVDGLDAKIIPGIASANVILKKVTDRYQVDLTDFRHLTRDVANVDLSHKQFLEVVTSQYPLMAEDYVCYPGGVSFSVKEKTTPLAEGKYLGHGVQKSVRYIENDGRSPCAAVALDLKKTVFHSEINALEYLKMQNTYVKQLIESKQRAKDNDMTLKTACQRIKGLRAYTQHTKRQRVVTMDRVGPETANEARFQLQDGEEITVAAYFHRTYGAPLLFPNAPVAVERKPGSKDVNYHPLELLTIAENQRVTGELPQAVISDVIKQAAVVPLDRQQQIMRAHRDLKISTNDYVDNVGTSIAPQMMQVKGRLLMTPKVVYGRNTQIEAREGKWRAERKTFLKPAGAARWTCMMLTNNRLGEQMMHNFLNKYVAVCRRNGMQMADPIEPFVVDWRRTDLQTEIDAFMKDCTQQYKLEFVLCIQDNNMHEHKYLKFLERQYGLITQDICTRTVERCIGNASATIDNIVQKTNVKLGGLNYGLEYKSPDGRHDVLSATTMFVGLGMSHSKPPKPDATGQEPSRPASVIGFAANVLAQSFAFVGDYYFQKADRDEKIFAIVPIVTRLLEQWCEHHGGQMPKNVVFFRNGGSEGQFQMILKYEVPLIKFAIEEFAKKSAVQQQFETKFCLLVANKRHNVRFFKSNITSGGRAAQQNLQPGVVVDSGVTHPVFSEFYVNSHTTLQGTGKTPRYTVLHNDCGLTLNQLEHFVFALSHGHQIVNLTTSLPTPAYIANDYAERGMAVLGQFLRASGAPVDEYETFNSKLTFAAMHPGSKFHHCRVNA
>WAGO-1 (R06C7.1) #11 - L596_g1696.t1
MERFKKLPPTTPLKARQVTLVTNHFRLNPTGRTVFRYDVAMSHTRMVKGTEKIRDLCKGDRDDAAILERNRRCLALMDAAFAAAPFASTSAAVIYDNSKTLIAAEELDMRQCACIRLEGGTIPPGFINHPRFKEGYFTIQITPTSSNHRLVINDLEAALAGDEANPDRSLRQFLEILTNQETLKKNSFMAFYGNLYSREAADQRKLREARILKSGMSKGARIIGSSSDLVAALVLDSKKATFFDDTNKNGLAGNIRELLNRAPNDAPSRVHINDFDRPDIVKYIKDLRVYCLNKPDNTFQISGLTREPLRNVFFDMGGEQLSVLEYHQRDGARLAYSHWPAVIVQSPRGRNYFPIEVLGVCEGQRVPISKQTPGQMKVVVNDCAVLPNVRFAEIHQMLNALNLASTTPNRYLQAFGVTIDVRPMKITAYRRQAPKIVYGGNIKVQYDDVKGSWFSSGPYVLPAKIPKWFVVYDGIDQRTVQQFVGVLQQAMKDKRMEVAQPKYMEMKVAGMDAFLSSIVKSLKPGERSPFVLFTDANEDSHAFLKLQEAKHQIVTQHLRTKTVRECIEPRKKLTVGNILNKLNCKNFGLNHLVAPDHEKNYLQKADVMVLGYDVSHPEPQSPQDRRLGIPPSTPSVVGFSFNGGQNPEMFIGDYQFCPPRQERVDILESRIQWMLKVFNDHRKKLPERIVVVRDGVSEGQLDMVLQHEMASIRRGAAMIKEGYKPKFLLVVATKRHQKRFFIDKSNGEVDNSMPLTVIDHTVVRPDVTEFFMQSHKAIKGTAKVPAYTVLQNELGMSLDEIQAFLMGLCFEHQIVSSPISIPEPVYQADQWASRGHSNVLAFFRVMDIEDPEDPSKKMINRYLKPVEDPLPGRPKFEYDWTRISKLLCYRGRRLEKTRANA
>WAGO-1 (R06C7.1) #12 - L596_g18749.t1
MPLITVEMSLFVAEGSVVIFFNAIVLLIIVTDSTLRESTELMFIGGLCFADMTDGIAYFYAGIHRLCNILSRTDDVMISRLECFQKPFMFFFFYGYQLPAMMIFIVALDRFVAVFAPMWQRKVERSSKLMVMVAIFLWVTLTYVINVFLFHSYSTGYTTAQCFAHDVFLSQLWDFIIVQRSILILLCVLLYVPILIKTRRIFQSNDKSVQSNSFNVTIGLTMTCSVLLLFIPDVIIYFDLIMDFHLILYLLGLNKCVANVFIYTLRQKEIRKKIELICRRVFCNLKGPFDLELSKRRQRSNITRTLGIAKRCQSLSAVAWSVVLSSAVPQETWHVISRPPLGEARFDGRIDKGFIAPLLYFYVSKMSAGPSNLQPDVELRPATKIRHAAEIKFKEMGIEVGIPEMPPKIPVPPNGNVKLTSNLYAVRTSSQIPIYRYDLDITISQRNGKQLALTKKSDSDAVSIDRKLKTKIIFFKMVQTYPELFSGVHHCIYDLESTLFTLNDIMKDDPEAEPKTLVIEGLDGAKFQGTTSATVILKKCHDRFEVDLTNFSHLELNIESTDLSHKQFLELVTSQIPMMSDSFVTFPGGVSFSMNASTTELPEGKYLGHGVQKSVRYVENPQTRKPMAAIALDLKKTAFHAVLTGLDYLSQIVDDRPLHNGQKLPNSVFGSFGALKKMKGLRCCLQYGNRYREVVIHEISNKTAQEHYFDREGQQITVENYFYQMYNLRLKCPTAPLAGEKKPGQKALCYYPLELIRVLDNQRVTGELPQKVIRDVIKHAAVVPALRIRQIQDSCADLGLFGNDYLQNVDTVIDSRPIGVEGRQLKNPKMVFGKNTSVESRNGQWPNRGAMRPPFFIPSTIRKWSVLMISNLQNGFASETMHTFVSSFINECRARGMTLPAPSDPWALNAREELKPQMEGFFSECPGLGIEFVLVLQDDCFHEHKFLKFIERKYNVISQDVNMKTVQKCLQRAAATLENIVQKTNIKLGGLNYSIEMTNPAGTSNVFQPDTMYVGLAMSHSKPPKPDSTGREPPKPASVVGFAANVLPQNFAFVGDYYFQAADRDEKIDSIVPIMHRILDMWCQHHEGQMPKNILFFRNGASEGQYKSILKFEVPLIKHALEKFRNNCGVEQMTPETKFSLVVATKRHNVRIFKANIQAGRPNEQNLPPGIAVDRSIVHPMFAEFFVNSHTTLQGTGKTPRYTIMHNGADFKIGQLEHIVFGLAHGHQIVNLCTSLPTPSYIAGDYADRGMVVLHEFLKTTRAEADQYDTFNEQLTYAALQPESRFHYCRVNA
>WAGO-1 (R06C7.1) #13 - L596_g18732.t1
MLGKWRTVRVEFEAAEELDFRWEFDHMVAAIFEALQPHCRCLLGFGTRSTFQINTVYPKELQSPAILFPDDPNGLCKRNLLVQVSLFLLLKSFQLHDFSSSQLSGRFLLLFRTCGESECERKRPRFATFGLPNLRSSANVQKLRYANSIAKTERAANRRSNPLQTNSSPGPARKTAPHRNPAANPKTCFAIGVRGQVADTVDRSFVPRGPLLTVAPPALSPSALPVFSPIRLFTPPHPAAYFRQSTMDLHTKLAELSVADNAMKPTPATEFKETVEVVSSYFEISIGQNALAYRYEVDILAISQRGQEKNLTRGPADDGAASLRRQLCHEVFNAALKKSNGFGTKQGVRLPVVYDCRATLFLPAPLKMEDEVVVILDKDADFAEMSKETLFTLDPTDVIRVRIAPTTQNAFEMDLRAELMKADFCEPDCEFARDRSFRTFLEMLTSVNAVRSGSHTQLGVGNFFDNDPSMIVDIGDAKCLRPGLSKGVRVVEKDDRPYPALVVDAKSSCFFKAQNLAQSIMELGRQKGRPDDMWKTARFLFKDVRVISAPVDKKGRVKSFPIRAITKMPATELNVKVKGFNGSLADYYARVLKIKLQHPNFMCVEADVPGPKKEFFPVEVLFVSPNQRVPIEKTEANQSSIVLKANAIKPDRRIKNIKDQMSRMSLFANTPVMEAFKVFVCPDFIRLTAGVRVAPQISMGDRDVKIDQKKANWAKEANGANYKQESFSLDSWAVLYANTEPGLIRQFVQRYVAAAQRRGFTVREPTILKFNQDFERTFSECIDNDIRFLMLIDPKYVKTHESLKLFERLCHVLTQHVSLERVFDVVQKNSRMTLDNILSKTHMKLGGLNYVPIIENVGSRFALDSGEVLVIGYDVAHPTAMSPQERRLVRSLNLDVKSLEPSVVGITANCSFNPHEFIGDYHYQTARKESIDVSILERRMVWIMRLLEKNRPDQCRPKHVVILRDGVSEGQYDMARNEEMAALRDGLKLVDPEYNPTFTLVIATKRHNKRFFGQDGRNYVNTDPGTVIDKTVVRKDVPEFFLQSHYPLQGTVKIPQYNKLHDEANFSMDELQAFVNCLCHTHQIVNLAVSIPEPIYQADELAKRGRNNYAELRRRFTSEVPRMNEAGVIDCNALTKILSYWDSPLEAVRFTA
>WAGO-1 (R06C7.1) #14 - L596_g20920.t1
MEPSSQPPPTPSLPENMAPKIVGPRDIYERPVPIASNLFPLEMKADVPIFMYHVQIHMKIGAHEINLVKRHTDDYMHIDHKDKCRAAFRFAVRSSPATFGDPKGLFYDLQGQLYSDHKLKDVLGHDLIVREEIGIPGEEAGRAPEFKNLGVEYLRVEVEPTRENRPTMILGEMMATRAKMQESVSRELTQFLEVATSQYAFLTPSKMVTYPSGVSYFKPTTAEPVQVFPAAKELLDGVHKVVKLIDGTKRGEMAVVVDPKKAVFHKSNITVIDKTVEMGFLDRGSGTVLRETIPELAKKLKNLYVETRYGKKPIRFAVHDVVLDTARTSRFNKNGDMTSVEEHFKDEYNVILKHPHAPLVVSTPLKKNPESLQLLYFPMELLFVCPNQRVLFNQQTAKEAFAIKKASTILPENRLKEVVNSASKLRINGASVQGCFKKAKIEVGNAPLTVEGRSLVPPHIEYRAQQVQVDAVSGRWRSSSRRGKPQYLVGGKIERWAMYVLSQAPSQAEEELGKNLLAKMEDEFQARGMQIAQAKFLATVKASPEYMKKIFDRAKAHQLEFLFFIQDTKICLHKEMKYYERKYEIISQDLNMETAKAVTEEGKYHALDSLIAKLNVKVGGTNYGLVGPSIPDLFKRGKLYIGFHASFSANPADAEDPTVIGSSANVTQTSAAFVGDMFFQEKNDVDMNTAMAKATLKYVLRYKNVHGHAPSEVVIYRSGSSEGQFGQILRDEVPALRNALQNAEADEAKLTLLMVNKQHSVRLMPSVIMPGSRAIEQNIKPGTVVDTKITHPRFAEFYLNSHQVLHGTAKTPKYTIIVDDSAHQIEYLERVTYALSYGHQIVDNPTNLPSPLYIAGEYADRGLTLIKAKRKLGDTVDVKDLEKDLPYMASQVLADKRVNA
>WAGO-1 (R06C7.1) #15 - L596_g16096.t1
MSNNERVKTIQNGMETISISATMPVKKKIANAPGAPLQVATNIYKVGLSQVPIFRYDVDITLKLPNGKDVKVVKKDRADHVIIQNKNKACKTFQKAVQKFPNVFGNVELFYDCQSILFSLRKLNISDKGQEFTLSPNDLPEVYGHVDSVNFIVKSVREKYQLTLNDLGFLSTQEVNGIKHDLAQFIEVATSQNALFNGKHSVYDGGVSYLMSTGEAVRNEPSKQLITGVQKSVRFIEGQGKPEATLVLDLAKTVFHKGGQNLYEKAISSVRNWPRNDQVDRREIKSLSQQFKGLSIKTEHQKTKEYELAGLSEFSAHTHKFDHNGKQMSVAEYFKKQYNKNLMHPNAPLAIAKNFFMGKRNTMMLPLEICTVLPNQKVSKQQETPLQTAAVIRYCAVLPSDRKEQIYTQVKELGFWGNKNLPITVDQQPIVVTGRQLPQPSIVFGGNQTVSVNPANGKWQATKGVNRNARLPFALPGNPMPWAVVLVGQGTQDTAKSFALAVKTECESRGLKMGDPSAVIQANYEDIEINARNNTFENLTKMTPRVKFCLVIENERSPAYIHALIKKNEQNWRIITQVTDSATVQQFNFALGPAQGYGKGQTLENICLKTNVKFGGLNHFIKPPQGHEKVFASDRMYVGLGISHASPISDAQRARNVKPSPSVIGISCNYLAHPQALAGTFVFQEPRENKMVESLKKVFFDLATKFKNIRKVIPKEVVIYRVGASEGQYATILEQEVPQIRAGLKEAGCNAKLVLIVPSRTHNVRLFPQTIKKEDKASFQNLKPGVVVDSVIVHPKFPEFFLNSHCTLQGTANTPRYNVLVDDLQAPISDHEMATYMWSFGHEIVGSPTSLPSPAYIADKYAERGRVLAVEARRKDEKFLNEEGEVDFTEMTAKLSFAGTPLANFRVNA
>WAGO-1 (R06C7.1) #16 - L596_g21757.t1
MSEISVDELSSYISLESMSIGSSSAYHKKAPVPDSSLHHPVELLSSYLEIGIQEGSKAYRYDVEIEASKSGSLTRGPADDGHGAMRRKVCYDLLYAALAKSKGFGTGQDYHMPLVYNRQNHVFFAQPLPEDFITIDLDDNADFTGITNYLYYMTGATESIQVKISKPQSEEHVLDLYESIFDTAPLDCDGNLEISSRDRSFRTFLELLTSGPASFTGSHVACGTSFYEATNGKDMKDGKTMRAGLKKGVCVIERDGILYPALVVDSKTGAFFKEQNLLKSMKEMNNGEVPRSAAEPMWQKARKLYKDVRVLVVSNVYKNSTQRRLTFPIQDFTREPASRQMMNMKGFRGTVEQYFRQKHNLSLLHPHLPCAIHASNSKIPVPTFFPIELLFVCGDQKVPLEKSDRFHSETLLRENAVDPKLRKERTEVQLKKLGLWREKREKMSVDDKKDLSEDGTNLLTAFGVSIINEFIPIRAGVRVAPTIEVANSEKISIVQKTANWEKKLTNKRYFSSVEITNWAVICSQISAENPMIVRFLKQMVNVSKRRGIRMDSPMKYSLKSGSREDFDDIFRHIAESGRHFVMYFSPLKEKQHDLVKWMEHHYSVVTQHVCLERIENVVTGRIQILDNIIHKANMKLGGLNVIPRIEKLGQRMEIESGDFLVIAYDVCHPAPMTSRERVLMRSMTSFDPSIRSLDPSVVGIVANCVAHPHAFVGDYHYQASRKESVDGRILVDRVKWIFELLAERRPNASRPKHIIILRDGVSEGQYKMAEHEELSAIRRAVAMIDPNYHPTFTLVIATKRHNKRFYDKNEGIAVNTEPGTIIDKDVVRGDVTEFFLQSHFPLKGTVKMPQYAILCDEADFSQDEIQAFVNCLCHSHQIVASAVSIPEPIYAADELAKRGSNNFAEHVKIHGRKSLRKDPMNPNLIDFEALTHELAYWKTNLEAIRFNA
>WAGO-1 (R06C7.1) #17 - L596_g18751.t1
MSDQPSSSHQLDVQLRPATIIRQDAEKMFKELGIEVPEPIMPARIPVPPNGNVRLVSNFYPAHVNGQVPIHRYDVEMTIARRNGSQLALTKKSTSDAVSIDRKAKTKAIFSKMVATHPELFTNMYSCIYDLESMLFTLKEIEEVNLIIDGLDGEMFQGTTSATVNLKKCHDRYELDLTNYGFLQADVGSVDLSHKQFLELITSQIPMMSEEFVCFPGGISFGYHVDATPLSDAKNLYHGVQKSIRYVENPATKKPMAAVAVDLRKTTFHQSINAFQYMCAQVQVDRNGCLTNASSLTKVSKAMKGIRCKLTYGRRYREMVIHAVVNKSPRSTFFKRGEQDISVEQYFYEVYQITLRYPNGLMAAEKKPGQRELCYFPMELLEILDNQRVSGELPQAVVSAVIKQAAVLPAQRRQEIQQACNDIGLFDNEYLRNIQTTVEPRPIEIEGRLITNPKLIYGNNVLVDSRKGQWPNRGANKPKFFLPAVVRKWTFLMISEQRNGQATGTMNGFLKEFIRECQMRGMTLPHPSSPYVLDTNRNVEDQLEEFMRDCSEGDYEFVFVLQDGIFKFHKLLKYLERKYQVITQDLKTQTGQRCLQRAAATLENIVQKTNMKLGGLNYSLQMTNPGGRKSVFDESTMFVGLSMTHGKPLKPDSTGELPPRPASVVGFAANTLPQQFAFIGDYYFQAADRDEKIDSIVPIMTILLTKWSKHHDGQMPMNVVIFRNGASEGQYKNVLRFEIPLVKYALEKFREAADVEQIHPETKLCMLVSNKHHSTRIFKTNVPVQGRAPEQNLEPGIAVDRAIVNPVFQEFYINSHTTLQGTGRVPRYAILNNDAGYALGHLEHIVFGLAHGHQIVNMTTSLPTPAYIASDYGDRGMVILTQFLREFEKLKGYEHPVDAYQEFNQSLTYASLDPDCKFNLCRVNA
>WAGO-1 (R06C7.1) #18 - L596_g4755.t1
MLSVEEQAIHPGRCGKRLQVNSNVLNVRLPASRIYHYHIDVVGHKRCGPKIVLSRSFFNDSLGHERRTVLVNLFNWLAYFNEERIFAQPRNFLFYDAKSNLYTRKKLNCKCGEVVTISTAQLTEAQIEGLEEFLCVRVKFTEADPFEIVLNEPEEFRDLGGPLQNFLETLFMQHSLFKNYESVVMNPLIQVAADYEARGIPEVEASTLVKDGSRIFPAIRTKLMNMEGPQALTMEGLHEADHACLCFRWEPFAVHSSVTLDQKARRALEKEGFESFFPFELDALNRELCGIRVFTELSGTKKYFIVDHVSERTAWTAVDGQTLKDFLKTEYEVTLKKPDLFLIVEKRGSLDIYHAMELCTVAPFQRPHKKLNVPPRFKWDCHQMKNKASPSKHTDHAKRLRESVGLVNENVFLQGSGVTLATEPMEVTARILESPMLRVKREGLFAVKSKGEWRYPPTTHFVHSAKIERWGVCLIFQERMWCSDRYQAHIRLNNLMKVISDRAGGHGMRMAERFQEPYNVPIREGSSLSDQIEQVRLCFERCKETFDPQFLFFVIQEGIHGLRDCIEAFERKYQIAALDVNNNEALKMALAHCSSRQKSPMAQVLTSTNIEAEKALRTLMAKINTKMGGLNFEVVPNHANRLLLQDGYLFIGIHAAFAWYEQQRATVVGYAANLRYPMAFSGDARVRKFDESLPEFLAKIVKRCCIQFNKVRGVDPQHVIIYQSQPSKDLCQLIVDRTAILLGEIGISTELTYVFVDEHHDIRLTNTHLNNTSVIEEQNLLPGTVIDTHLVRKGASEFFLNSHIGLVGLSEVPKYTAHDFRSGLDGDNLQSLTYTLCYAVQSLNAPVSVPAPLYVAKNKARHGASCLKLTGKVGARDFVN
>WAGO-1 (R06C7.1) #19 - L596_g20173.t1
MADEVIGRMRQLSICDDPAYALAPKLQRADKKSFVQHVDIVTSYVRINIVGPAKSYRYEVTIEALAEGREPNILTKGPADDGHAVMRRTVCYFLLYAALDKSGAFGTGLGHELPLVYNGQTLVYFAKPLDRDVIQVTLDQHDDFANVPEELLYMVSGNDTVRISLQKALVDSEMDLQTQMFETGQADVENLSTHDRSFRTFLEMATQQAAHYAQSYTSIGNAFFEKDPSRTRSLGDGKIIRPGLTKGVRLIERDGVVYPALVVDARSAAFYKEQNLFLSVKECMDSGDPNMSMNDKWDRIDNLFRGMSIFLAT
>WAGO-10 (T22H9.3) #1 - L596_g19923.t1
MSFRIPKKRKPEDEGAGGTIPPKQNYSSSSSYASASSSSRQTSNGGKNASVYMNGFELQIGRSVEIHKYEIKLFGVFRKRNGDEDRKDLTQGSKEAKDDISIQKRRRGCWEIFRAVVRENNSLFGDRNHRFVYDCGLIFYSIDAIFPETETKTGTFDVAILSRECQDYYGQAMKMIEYSIKKVRDGTFTLGVAPEDKDDRSMQQFLEVLTSQGVYAEGLDNLIFRNHRYDVKYDTPVLRNKPEYPFCVRHGSSKSVFVAEIENSNKALQTILQLEKKTSPFFPRMNVLQFVDQSKSDIGERIVRGLFVTTTHLKKQKVFRVADFSKMNCNDITFKMRDRQNEDEFREISVTEYYQEAHKFSTRAGHRPCVVERKKRFNGEKEENNYPMDCLEIMDGQRIMDKKQSGDITAYLISEARVLPRQMGDEIKEELNHRVLNQEAEKYLEAFGVRISRDLLRSDAKILQAPKIAYGDKEKYLTTDNGQKNAWKIDDALKFYRPGKISDAGGENEQNRWIFAVLNEFQSERDHAAAKRFLGKLQDRAKLRGMLMLNPEVRCLDVRDPDPDTVNKKLSELCQYAKANKVKFIMFIQYERKDMSRDTMKQLETKFKLTTQQITMSTVNKGAGDKGDRMVLDNILNKTNEKLGGINCIMKPSPQIAQWFSGNVMYMGLDISHPGLGGNALSSVTPTAVGMTFTKNRDEVQGRYWFQEAREHMLRSLKKQIVFAIEEFNRCSRRYPDKIVVYRGGVSEGEYDKVKTDEVEQFMEAFREINFPGKRKPALIIVIVQRNSGYRLIPTQDNDFRGNDAIVQNVLPGTCAEVIGEGGRKEFILVPHQAIQGTAKPSKYVLIYDEAKCITLSELETLTNTLCYSHGIVTSPVSCPSILYQAGDLAKRAINNYRIHSSRRDFGAMPPIEEVDKRNEYFDQMCDMLQVTLDTRFWA
>WAGO-10 (T22H9.3) #2 - L596_g15911.t1
MLPNRAPQGNELQRNRKVELQVNGYRMRINESKVYMHEIKMEVAFNTLKGLRNVDLLARPANDVIRQKRRRLIWSIFHATRIKHPQQFLYNEYEYVYDCGGALFALHQIGDGNRIEFVMKMADFPEEAKGQLHRAEHVILRLAFTRILDLRQSNLYDGGEAGEQRCRFVQQFLDILTSQYVLGSDQHLVFQNSRYSADPREDIEAHLSKVIKKGASKTINIVGDQKNQEALLFVEPRRSPFMADKKVLDIVEEVRRELGGRSSNAQLKKKLEDLLKGIVVETIHQKDAVQFPVKGFSAEPAGVLSFTMDAGESTTVADYFKRRYHLHVDRDMLCVVCERRQQKFYFPCEVLVVLPGQRVHCSRQTPKLVEQLIKESQQLPSRMKEEVNHEREVYGFHELNQQLKAFNVTVDTELCTAVGKVLPPPVIQYEQRTVSADVDRSGQKITGRQWKVSGQRFVRPAPTPEKWVLCVFENALESESTRTFARAYVNAARSHGLILGEPIIERLQEVNQSTIYARGQAYKSNDVKFILFIFGGDRKNFERDIMKESETLFNYTTQAVNTKTAMKAISDRGAFMVLDNLVMKTNLKLGGINHELANAQDFPQGYLEKVLFKAGRVFIGLDLQSPGMLGGADEFTLDPTVVGMTFTLGSPADMRGTYWYQPAKKKYISRLKDAIEDVLMVYQESVGSLPNDIVIYRAGVSEGDILMVVSEEIPAIKDYLTTLDNPDGCPYRPHLTVMVAQKNSTMRLMPLVVHQTGRAQDENVEPGTSVSTNIVSARHTEFVLAAQQALKGTARPTRYIVVHEEEGQFTVEQLENMTHQLCYLHGIVALPVSRPSPLYSATDLVKRGRANWKVRMERKEGSHTPLGPVNDTFFDPINEERLRFLPVTLPAKFWA
>WAGO-11 (Y49F6A.1) #1 - L596_g17524.t1
MNKSSKIAAMPSFKKKRPAENDDSPAPKRPAEKARRSQREEAPRRSHGGSKILMNGFRVEIEKAMTIHKYYIQLNGIFKKRGGEEIARDLTEGVHRDDVSMQRKRRGCWDVFRQIVKENESLFGGNTHKFVYDCGRLFCSMEEIFPKSETKTGTVDLATLPDRSQTFYRGALRIEWTIKKVEDGTFKLGVPEEARSSDRSAQQFLEILTSQGLYARGDDHLIFRNQRYDVQENAPVDKKSTYPFCVRQGVQKSVILCESAEKIATILQLEKKTSPFFPEMNLLDFVRQCSSDDKAMAVLKGLQVQTTHLKRQQKTFRISGYSNTACKDIHFENRDGEKISVPEYFLKHHAFRTAAGQLPCVEEKRKPQNNHYPMDCLKIVGGQRILSQKQEPEIVEHLISTARILPLKMADEIGEQLGKHILNREAETFLRTFKVNVDRNLLETEATLIKAPQIQYGKNQNGVNPAKTLTTERGQKNAWKMEDSLTFYRPGVVSDSGEAHRWMFVILNEPNQREHGDCRVFLEKFVARAGTRGIKIQFPHVEKKRIENSDPWPELQEIGQYAKSNGVKFVMFVYERTGDIRKAMKLLETTFALTTQHVSLKTISKAAGDKGAFMVLDNLLMKTNEKLGGLNTRVKAEPRVAEWFERGTTMFMGFDVSHPGLGARNENGVTPTAVGMSFTKNSDLEVVGRLWYQEPREHLIPNMKDHIVEALETFKKHSGKFPELVVVFRGGVSEGEYEKVQTKEVQEFQDAFQQLKFSRKPILKIVVVQRNSGYRLMPAQRNDFGYQKNEALVQNVVPGTCADAEIVDQKRTEFVLVPHQAIQGTAKPSKYVLLHDEAPKMSKEELSTIAHTLCFMHGIVTSPVSCPSILYQAGDLAKRASCNFKAFLNRKGGNVPIPPVEEKEKRKEFFDALCEKLKITLDTRFWA
>WAGO-11 (Y49F6A.1) #2 - L596_g17422.t1
MPFPRGKGYHPYRGKSYHRGFGRNNFQYRQTKREEYHSSDSSPTDNQPSTSRSHDRCDDYKPKDEDDMKLYLSDRKPSPEPPEFQIRINAFPFNIDHAPKEVHTYELIFVMSQKLKKEKEPANHKLRTNAKLFSAEDIGLGWFDAEEEGQVYHGTDMSCGPTDVVRRQRRKALLFQLFRHLINQSKEYFPGSKYVYAYDGQRILYSPEVLKMDEGMFMAQLTDLPESVTQLLGPSDQENTEINAYIRKAEDVIDLHDFGTEARPIHGIEGFLETLTWQHAFEGFEEHIVYGTRYFALNAGKKLKHCTGFKSIVGFEKRIELLPDYTCSNILLERSLRPVMRMTPKFDLFFDNSTAISLDDFAVLFFRVEVDDLSATLQKKENLQKLNQVFKNAVVRTIHRDDGRQDTFMIDHLDERNPFEITLKEEDEETVADYLYGVYGYKVDRNDRLPCVARKFKNEFAYYPLRTLVLLPNQKVASKFLPEDMRKSYQEACQNLPSDALNNIFTAMSQLNLSRASEDYNNDEKVIHVNNKYMENFKITLESNQLIQMPAQRICEPRIAYRESTHLAQDGAWDYEKAAVFIDAVIKVRRIGLVNTCEDVGEDDLGEFISKLIGWLRNKTIDLKLTPEDIFWWPDAWNEFKAGKSKKEMLATAEKIIETSEVKLMFVICSGSADDEIHDIWKLAEVTNELVKVNKKDFVTTQCITPATLSNVLVKSTYQDEILTNIIMKMNLKLGGTNYVLSKSPSNNEMSIPHIHSSRMFVGIDVLQPEKTQLNGEPTNNPTVVGISFTDATSKFYLRGTYWYQQSPTVSLCLLQKHFDEALYWYDCHKMDSVPREVFVYWRDSRIQKNMEEVKMVLEDVICNRAKDVRRDASKLFLIMVDTKPKTRLFTWETQFSGNAQTQNVQAGTFVRESYRKRQFTMINHKSGAGLAHPVRFTMINDDVNEKDYVEAELEKTTNALCFLQNTSTRSTSIPAPLYSAMDLAKRGMKNYETMDAVMREEERDEDRKKREARTPEAWHRYYKQLVKTHMSVMPIRDSKFWA
>WAGO-11 (Y49F6A.1) #3 - L59 6_g18655.t1
MSPPRNGSNFNRGNGHKRPPHGHNSQAKRVKWEEHSSDSSPIYNQPSTSRSYDRQPKEENDVKPLFSSRERSPRLRDLQIRINAFPFNLDHAPKEVYTYELIFVMSQKLKREKEPENYKIRTNAKLLEAEDVGLGWFEAEEPGQFYHGTDMSCGPMDVVRRQKRKALLFQLFRHLINQSKEYFPGSKYVYAYDGQRILYSPEVLKMDEGMFMAQLADLPESVTQFLGLDDQKNTEINAYIKKSDEVIDLHDFGTESKPIHGIEGFLETLTWQHAFEGFDKHIVYGTRDFALNAGKKLKHCTGFKSIVGFEKKIELLPDYTSSNMILERRIHPVMRMTPKFDLFFDNSTAISLEDFAVLFFRVEVENLPATLQKEENLQKLNQVFKNAVVRTNHRDDGRQDTFMIDHLDKRNPFEIIVTEEDQETVADYLYDVYSYKVDRNDRLPCVARRFRNELAYYPLRTLVLLPNQKVASKFLSEDMRKSFQEACQNLPSDALNNIFTAMSQLNLSRASEDYNNDEEVTHVNNEYMENFKITLESNQLIQISANRVHEPQIAYQKSVKPAQDGAWAYEKGAFFVHPKKGVRKIGLVNTCEDVKEDVLGEFISKLIGWLRNKDIDLKLTKEDIFWWPDAFSDFGVMKSVKEMLAKAEKILETSAVNHMIVICNGSAEDKTHDVWKLAEVTNGLAKKDRTNFVTTQCITPTTLGSILVKSTYQDEILTSVIMKMNLKLGGTNYVLTNSNRNYMSVPHITQDRMFVGIAVLQPEKTRLNGDATYNPTVVGLSYSEGSPNFYLRGTYWYQQSPSVDLSILKKTFAEALYRFDKFHLIPKEIFVFWRSGKFQRSMEEEKIALQEVIDKRVKDVNAKSPKLFIISVNTKPKTRFFTWETKFSGNAQTQNVQAGTFVQESFLRREFTMINHKSQAGLAHPVRFTMLNDEIGEQDYAEAEIERTTNALCFLQNNSTRSTAVPAPLYSAMDLAKRGMKNYETMDAVIREEESEEERKKRETRTPEGWQKYYGNLMERHMPVPIKSSKFWA
>WAGO-2 (F55A12.1) - L596_g16917.t1
MHAHRLQQLTDGINRNLNDEGIARRTPMSPTPSVGRHDKEITLTSNLYELHLAHGKVRVYHYTIKLSDYTGKTAEVHGDYGRVNLGSKLLLAADTFMKRNGFRDHDFFCDLAGMRLYTIKPIPRSEGNPRFYVSEATKDGNFAVCRDDGRPFELNIRDVFKEVRIPNLQDATLDAFINTAINKACLAKSDLYFPIFETCYPKTGAPVARKDGRSLILGTKTSVERVEGRYGSSTTPVVALNVSAEWFYMKKNLLDFCKQNIELNEKSVPVDAASFQLLSDSMQGVTVRLICSKNPLLFTIKSLANKNVSMLKYPLPNIKGTGLIKFLWETYSVRLTYPESFAAEVKPDFHVTDPRPIFYPVELLEIMPMQRALKGKKSCDGSAVDREKKIQEKVHQMERHVQKAGLGLSMDDAPVEVEAGVLDLPKIVFADEKVVEVNPSSASWRFGSGECPERFARPAEVKELPWCVMLVSDVPPTKGMSKKAKFFADLLKKQAAERGLQMKEPMYHPTKQGKVEIERFFNTTAVEFFVFLMAKSLDYHDFTKILERKYQVITQTVKMENAFDVVEDPNSTKSRKIVEHIVMKMNLKLGGVNSTVKPSRHLFSHEEVRRLLIGFTLIEGPKITGRDAAAFNRFGGKIPAVVGFSANMGSLEHEFLGDFTFQYLDHTNIVQDIKKIVATILERFRKTRRGRDPAEVVIYRKLDEFHFPRVIQNEIAPLKNLLEEKCHGFVSLIYIAMTKTHNIRFFPKGPPPEGDERKPNLVPGTVIDSGAVGGGLKQFFIASYSASTGTTRPPRFTILENSKEAKIRDLEKLTLELTFAHQTSTRSLGVPAPLVVAKNYARRGNVLAKNEAEKISDVDTSVGVETMIDRLPYADAPVLRDKRVTA
>WAGO-5 (ZK1248.7) #1 - L596_g12936.t1
MEGEGTELRPSTVVRQARNAAASSAGAPPAAPGSRAATASVEKSFYSKLVNVPTPPPEEKKPHGTAGRQLKLRTNVYGLSLPKDVQVFRYSVDASGTLQRNDLRIEFAKRVANDITYLNRREKCRRVIDQVVAKYSAIFGDRRELFWYDSQSILFSRNQLDISSEGQFVLDQSDIGQNPLFEGFAHLKMVIRPAQTNFAVSIGDLEAYIQAEVFESDHALQQFLEILTAQYAFNTPLEAMSFGSRTAYLLNPEKYGFKPADCADVGDGKFLGVGCDKSVRFIEGPGGAGSQRAALVVDLKKTAFHKDQSLYEKAREILNNRDPKSTDASRLRMQLKGIVVETKHGSRRQEFAVDNVVADTPATKKFKDLTGQEVTLQQYFQQKYNITLQHPDSPIVLTDRTKKFAAFPMEVCSVVDGQRVTLAQQTPVQIQKMIRQCAVPPADRQRQILGLVQGLQLNSENKYHKAASVGITPTALQVQARLLQNPTIVYGRNSTMKPDEKATWRLARQKPVYLKPAKVDKWAMFVICGGNRSDCVDQDILNQFSNMMVQECRARGMTVSDPTGFSFIGASREVVQETLEKAKTEGNQFCFFITNNDVTNIHQFMKFQERKLSIVTQDMKMSSAFDVVRKGKRQTLENVVNKTNMKNGGVNYSLRFDDPAFSMEKLLPKDRLVIGLATTHPKPIVGKKEQDEGPHDKKKQMHQQRTGPPVPSVVGVAANALTESIEIVGDCLFQQPNREEKIALLQPVIRSLMLQFMKHRGMPPAEIVVYRQGTSEGQFRDVMELEYKMVKAAALQQGLNPKITFIVVQKMHNVRLMPMDSKAGDKAPEQNVKPGTVVDTMVTHPKYNEFFLNSHVALQGSARTPRYTVLYDENRLPMDEIEALSHSLAFGHQIVNLTTSLPAPLYIANRYAERGHNIFIASQEDYTKSKTSFQSPHSTTIEGNLDFGRMMNELSYCNSELKDKRVNA
>WAGO-5 (ZK1248.7) #2 - L596_g12936.t1
MEGEGTELRPSTVVRQARNAAASSAGAPPAAPGSRAATASVEKSFYSKLVNVPTPPPEEKKPHGTAGRQLKLRTNVYGLSLPKDVQVFRYSVDASGTLQRNDLRIEFAKRVANDITYLNRREKCRRVIDQVVAKYSAIFGDRRELFWYDSQSILFSRNQLDISSEGQFVLDQSDIGQNPLFEGFAHLKMVIRPAQTNFAVSIGDLEAYIQAEVFESDHALQQFLEILTAQYAFNTPLEAMSFGSRTAYLLNPEKYGFKPADCADVGDGKFLGVGCDKSVRFIEGPGGAGSQRAALVVDLKKTAFHKDQSLYEKAREILNNRDPKSTDASRLRMQLKGIVVETKHGSRRQEFAVDNVVADTPATKKFKDLTGQEVTLQQYFQQKYNITLQHPDSPIVLTDRTKKFAAFPMEVCSVVDGQRVTLAQQTPVQIQKMIRQCAVPPADRQRQILGLVQGLQLNSENKYHKAASVGITPTALQVQARLLQNPTIVYGRNSTMKPDEKATWRLARQKPVYLKPAKVDKWAMFVICGGNRSDCVDQDILNQFSNMMVQECRARGMTVSDPTGFSFIGASREVVQETLEKAKTEGNQFCFFITNNDVTNIHQFMKFQERKLSIVTQDMKMSSAFDVVRKGKRQTLENVVNKTNMKNGGVNYSLRFDDPAFSMEKLLPKDRLVIGLATTHPKPIVGKKEQDEGPHDKKKQMHQQRTGPPVPSVVGVAANALTESIEIVGDCLFQQPNREEKIALLQPVIRSLMLQFMKHRGMPPAEIVVYRQGTSEGQFRDVMELEYKMVKAAALQQGLNPKITFIVVQKMHNVRLMPMDSKAGDKAPEQNVKPGTVVDTMVTHPKYNEFFLNSHVALQGSARTPRYTVLYDENRLPMDEIEALSHSLAFGHQIVNLTTSLPAPLYIANRYAERGHNIFIASQEDYTKSKTSFQSPHSTTIEGNLDFGRMMNELSYCNSELKDKRVNA
>WAGO-5 (ZK1248.7) #3 - L596_g17875.t1
MVCKGQATSEIELTTNAYGFLPFLSAKVFQYDVEIVGILSGTGRTVNFTKCSKHDAFRAARIEECRDLFEMVKQKYPEVFNAPDSNYFYDNGRRLFTKESLLPPLVSKQEFPLNEFDTVYLDRDYSRFDKILFSVEKAAEDPMDVKEVLKHVKDSGKSIRLNQFLNVLTSQHALSNPVKFATYRIGSAFFINADRYDHNHGVEDLTDDKEIRVGCEKSVKLIAGSSDDGSAIALVDIKRTAFHKSGGTLLKKAREVLGKWPQPCDANRLKPHFIDLAVYTQHGSKDRRYVIDDVIAETPATLKFAWVRGSKELTLVEYFHQAYSIEIKFPRTPLAVAMAKEGRTIYLPLELCFVSPHQRVTTAQQQATEEMIKRCSIPPLDRQKRIGRIVEAMKISDNPFLQIADTGCKLLSTIPLSVTGRVIAPPKIRYGHSEIQKSAFLKPAKINSWAIVVLTAQDDPELARGDILSPSVLTKFARLFRKECKARGMQLPDPFLKEFMKADVDQLGELMRCLAQGDSTECRPPLRFLIFVTNEKLTDLHHPMKYFERQCGIITQDMKMQTVVDVVLHKKRQILENIVSKANIKNGGLNYSVIIPDVPGKRPILGSGRLIVGLFTSPTFKWQFEDSPSRPTALGYAANTTPNEGEFIGDCLIQKGRASAIQTILRRVLAEFKKQRRCDPADVIIYRSCEEGREKKTLEEDLAAVRSILKSSNPMPTLTVIAVQKRHGLRLMPTAIQRQGEPQSENLKPGTVLDSCLTDPALTEFYLNSHATLQGTARTPKYTVVHNDVGLSLEEMETLTYALSFSHQIVPLPTSLPSPLYIAGTYAERGVSLYQQDRERGREGSLESCFTYGSSPGLKHLRITA
>WAGO-5 (ZK1248.7) #4 - L596_g24399.t1
MDVLTEAMSKMLPMNIAPKIVGAQDPYERVVPLTANMFPLHMRAEVPIFMYNVQVFMKVGFREVNLVKRNTDDFTIIDHKNKCRSAFRFAVRAAPQVFGHPSGLFYDIQAQLYSVRELKDVLGNDLKKKEEIIVPGEDARKAYDFQDIDLEYLRLVIEPVNGTNPSINLGELVLKQKNFSDEVPCELLQFLDVACSQHAFLTPTKFTTYPGGFAYFNPTSEEPARELPDAARLHNGVHKSVKVIEGSCTAGRCGELAVVLDPKKAAFHKPDITVVQKIQEMGFLQMASENVAPHRIPELAEALKNVFVETRHGKRRSRFAIHSVVAESARTNRFTKDDGQVTVEEHFKKEYDIALKYPHLPLVVSMPLRKKTPSNGRAPPRLLFFPMEVLFICPNQRVLRNQQSAKQNNEVIKSCAVAPEHRLKDVIASGQKMRINGPNVHGCLNSAQIQVESEPLKVEGRTLVPPNIDYKGCQVQVDSFTGKWRNFGRNKPHYLEGGKIGRWGLYVLSKAPSSEEEQLALKFKDKMLVEFQSRGMQIDLPMFLATVKATPLYLRTIFEKARKERLEFLFFIQDKDLALHNEMKFYERAYEVITQDLRTDTARAVIEQGKNLSLENIIAKLNVKVGGTNYSVNGPSVPDLFKKGRLYIGLQASTNGPPAAGAHLPTVVGSAANVTTAPSSFVGDIYFQKFGEMDLQGAMASTTEGYVKRYAAVHGRAPDEVFIYRSGTANTNIGQMLRDEVPAIRCALKNSGASRARLTLVMVTKQHNVRLMPTNMTLGGRAIDQNIKPGTVVDQKITHPRFAEFYLNSHQALHGSAKTPKYVVVADDCSNPIQYLERVTYALSYGHQIVGMPTSLPSPVYIAGKYAERGAALLQTKRNLGGALDVDALAEELAYANSKVLGFKRINA
~~~

